# Investigating Protein Degradability through Site-Specific Ubiquitin Ligase Recruitment

**DOI:** 10.1101/2024.11.11.623099

**Authors:** Olivia Shade, Amy Ryan, Gabriella Belsito, Alexander Deiters

## Abstract

We report targeted protein degradation through the site-specific recruitment of native ubiquitin ligases to a protein of interest via conjugation of E3 ligase ligands. Direct comparison of degradation ability of proteins displaying the corresponding bioconjugation handle at different regions of protein surfaces was explored. We demonstrate the benefit of proximal lysine residues and investigate flexibility in linker length for the design of optimal degraders. Two proteins without known small molecule ligands, EGFP and DUSP6, were differentially degraded when modified at different locations on their protein surfaces. Further, the cereblon-mediated degradation of the known PROTAC target ERRα was improved through the recruitment of the E3 ligase to regions different from the known ligand binding site. This new methodology will provide insight into overall protein degradability, even in the absence of a known small molecule ligand and inform the process of new ligand and PROTAC development to achieve optimal protein degradation. Furthermore, this approach represents a new, small molecule-based conditional OFF switch of protein function with complete genetic specificity. Importantly, the protein of interest is only modified with a minimal surface modification (< 200 Da) and does not require any protein domain fusions.

## Introduction

Targeted protein degradation, or the use of small molecules to redirect native degradation machinery toward defined targets, allows for rapid and reversible protein knockdown.^1^ The most commonly applied tools for targeted protein degradation are proteolysis targeting chimeras (PROTACs, Fig. 1A),^2^ which are heterobifunctional small molecule degraders containing a ligand (green star) specific to a protein of interest (POI, orange) tethered via a chemical linker to an E3 ligase ligand (purple triangle).^3,4^ This design induces recruitment of a native E3 ligase to the POI, thus promoting the ubiquitination and subsequent proteasomal degradation of the target via the Ubiquitin Proteasome System (UPS).^3,5,6^ The ligand for the POI is typically derived from a known, high affinity, small molecule binder.^7^ By exploiting the UPS, PROTACs provide several advantages over classical small molecule inhibitors, such as lessened restraints of POI binding location, as protein function does not need to be blocked, catalytic function, and the prolonged response of degradation as compared to inhibition alone, thus expanding the realm of “druggable” proteins.^4,7^

**Fig 1.**
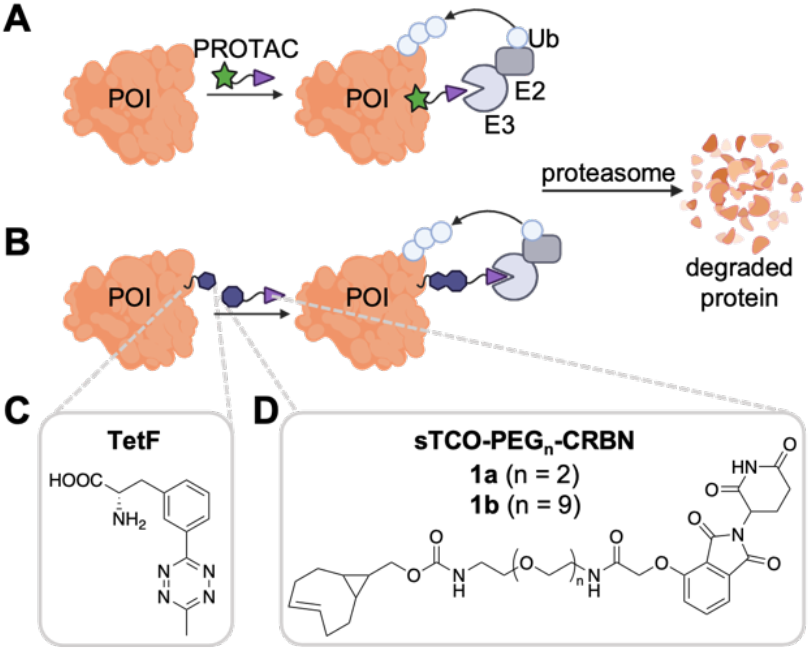
A) Mechanism of action of PROTACs. B) Site-specific conjugation of an E3 ligase ligand for targeted protein degradation. C) TetF amino acid site-specifically installed at defined positions of the POI surface in human cells with an expanded genetic code. D) Cereblon (CRBN) E3 ligase ligand modified with an sTCO handle for tetrazine ligation via an inverse-electron demand Diels-Alder reaction.

While classical PROTACs show promise toward previously “undruggable” targets, their successful design requires prior knowledge of a small molecule ligand specific to the POI. PROTACs are further limited by the generality of the binding location of the known ligand, as the ligand pocket may not be proximal to surface lysine residues necessary for ubiquitination and subsequent degradation via the UPS.

Some targeted protein degradation methods have been developed for proteins without known ligands, including the degradation tag (dTAG) system,^8,9^ Halo-PROTACS,^10^ or the auxin-inducible degron (AID).^11^ However, each require the generation of protein fusion constructs, thus potentially impacting the native folding and function of a POI and not providing information on binding site selection for small molecule PROTACs. Additionally, inverted approaches have been used to investigate the ability of a library of E3 ligases to degrade a protein of interest. Specifically, functionalization of JQ1, a prominent ligand of BRD4, with a maleimide moiety allowed for conjugation to recombinant E3 ligases. The function of these ligase conjugates required cells to be permeabilized for protein delivery.^12^

As the scope of PROTACs continues to grow, a generalizable method for bottom-up experimental evaluation of PROTAC binding sites, linkers, and E3 ligase ligands would be invaluable toward the development of novel degraders for any POI. Thus, we developed a tool that allows for the precise, site-specific recruitment of an E3 ligase ligand to any protein of interest – even those without an established small molecule ligand – without the need for protein fusion domains.

Utilizing unnatural amino acid (UAA) mutagenesis to infer bioorthogonal reactivity to a protein surface, our approach can elucidate optimal locations for PROTAC binding based on the surface properties of any protein. We hypothesize that this tool will allow^1^ for (1) targeted degradation of a POI without any known small molecule ligands and (2) elucidation of optimal protein surface microenvironments for improving classical PROTAC designs.

Numerous approaches exist to modify native proteins *in cellulo*, however, the only way to ensure precise site-selectivity across a protein surface is via UAA mutagenesis. Genetic code expansion utilizes an aminoacyl-tRNA synthetase and tRNACUA pair evolved from orthogonal systems. The pyrolysyl-tRNA synthetase derived from *M. barkeri, M. mazei*, or *M. alvus* is favored for its orthogonality in both eukaryotic and bacterial cells.^13–15^ The synthetase/tRNA pair then encodes the desired UAA in response to an amber stop codon (UAG),^16^ and the pair has been engineered to encode hundreds of UAAs.^17,18^ Azide-containing UAAs, derivatives that include azido-phenylalanine,^19^ azido-lysine,^20,21^ and azidobenzyloxycarbonyl-lysine^22,23^ have been utilized for Staudinger ligations to phosphines,^22,24–27^ photo-cross-linking,^28,29^ and strain-promoted azide-alkyne cycloadditions (SPAAC).^26,30^ Similarly, strained-alkyne UAAs including cyclooctyne lysine,^31,32^ and bicyclo[6.1.0]nonyne lysine (BCNK)^20,33^ have been reacted with azides for SPAAC. The fastest protein bioconjugation reactions are inverse electron-demand Diels-Alder (IEDDA) cycloadditions between tetrazines and dienophiles.^34–36^ UAAs functionalized with dienophiles include analogs of *trans*-cyclooctene lysine (TCOK),^33,37–39^ norbornene lysine (NorK),^40,41^ cyclopropene-lysine (CpK)^42,43^, and BCNK.^20,33^ Tetrazine-modified UAAs include several phenylalanine derivatives.^34,44–46^ The fast kinetics, small reactive handles, minimal toxicity, and superior efficiency of IEDDA reactions have made this method a favored approach for *in cellulo* bioconjugations.^47^

To combine the advantages of site specificity afforded from genetic code expansion with the potency of PROTACs for targeted degradation, we have designed a system that utilizes a bioorthogonal E3 ligase ligand for the site-specific, completely selective degradation of proteins (Fig. 1B). Our bioorthogonal E3 ligase ligand consists of an IEDDA-reactive handle for reaction with its UAA partner displayed on a protein surface, connected to an E3 ligase ligand via a polyethylene glycol (PEG) linker. The two E3 ligase ligands most commonly used in PROTAC design are the thalidomide derivative recruiters of the cereblon (CRBN) E3 ligase^48–50^ and the peptidomimetic ligand known to recruit the Von-Hippel Lindau (VHL) E3 ligase.^7,48,51^ Additional small molecule ligands have been used to recruit MDM2 and IAP E3 ligases, but are less prevalent to date.^48,52^ As CRBN recruiters have been used in the majority of reported PROTACs,^51^ it was selected for use in our degradation studies. Further, many PROTAC linkers have been employed, as linker composition plays a critical role in PROTAC conformation and activity.^53–56^ As no standard method for linker selection exists, we chose PEG chemistry as an initial starting point due to its prevalence in therapeutics. As this methodology expands, we aim to further expand the linker compositions for expanded screening of degrader designs.

We recently developed a method for the quantification of bioconjugation reactions with UAA-bearing proteins expressed in live cells, which is essential for the methodology described here.^57^ Briefly, a chloroalkane (CA) carrying a bioorthogonal reaction handle is utilized for conjugation to the selected UAA. We can then quantify the cellular protein labeling reaction through a simple western blot of the HaloTag conjugate (Fig. S1A). We found that tetrazine phenylalanine (TetF, Fig. 1C) was the most amenable UAA for incorporation in mammalian cells due to its stability. Excellent labeling of TetF with sTCO-CA occurred in under 60 minutes with concentrations as low as 10 µM. However, we found that the bioconjugation efficiency was protein-and site-dependent.

## Results and Discussion

With our previous success in labeling and quantifying *in cellulo* bioconjugation to proteins containing TetF,^57^ we utilized the same chemistry here. mCherry-EGFP-Y151TetF displayed near complete bioconjugation to sTCO-CA, which prompted us to use this reporter in initial studies of protein degradation through conjugation of E3 ligase ligands consisting of a strained *trans*-cyclooctene (sTCO) linked via PEGn groups (n = 2 or 9) to a CRBN ligand (see **1** in Fig. 1D). To first establish an optimal concentration for degradation by **1a**, we incubated cells expressing mCherry-EGFP-Y151TetF-HA with increasing concentrations of the small molecule, keeping the compound-containing media on the cells for the entirety of the experiment. For overnight treatment with **1a**, optimal degradation was observed between 1 and 10 µM (Fig. 2A). Unsurprisingly, a distinct “hook effect” was observed at higher concentrations of **1a** (100 to 250 µM), indicating saturation of CRBN with non-conjugated ligand, thus preventing degradation of the bioconjugated protein (Fig. 2A). Further, incubation with **1a** in the presence of proteasome inhibitors prevented degradation of mCherry-EGFP-Y151TetF, validating that the observed degradation is mediated by the UPS (Fig. S2A). Further, a competition experiment in which cells expressing mCherry-EGFP-Y151TetF were pretreated with a thalidomide derivative (Thal-OH) prior to addition of **1a** demonstrated inhibition of degradation, supporting cereblon-mediated degradation (Fig. S2B). We have additionally validated the mechanism of **1a**-mediated degradation through siRNA knockdown of cereblon.^58^ Co-transfection of cells with CRBN-siRNA and the required genetic code expansion machinery reduced the amount of mCherry-EGFP-Y151TetF-HA degradation by **1a** (Fig. S2C) in direct correlation to the decreased amount of cereblon present in cells (Fig. S2D).

**Fig 2.**
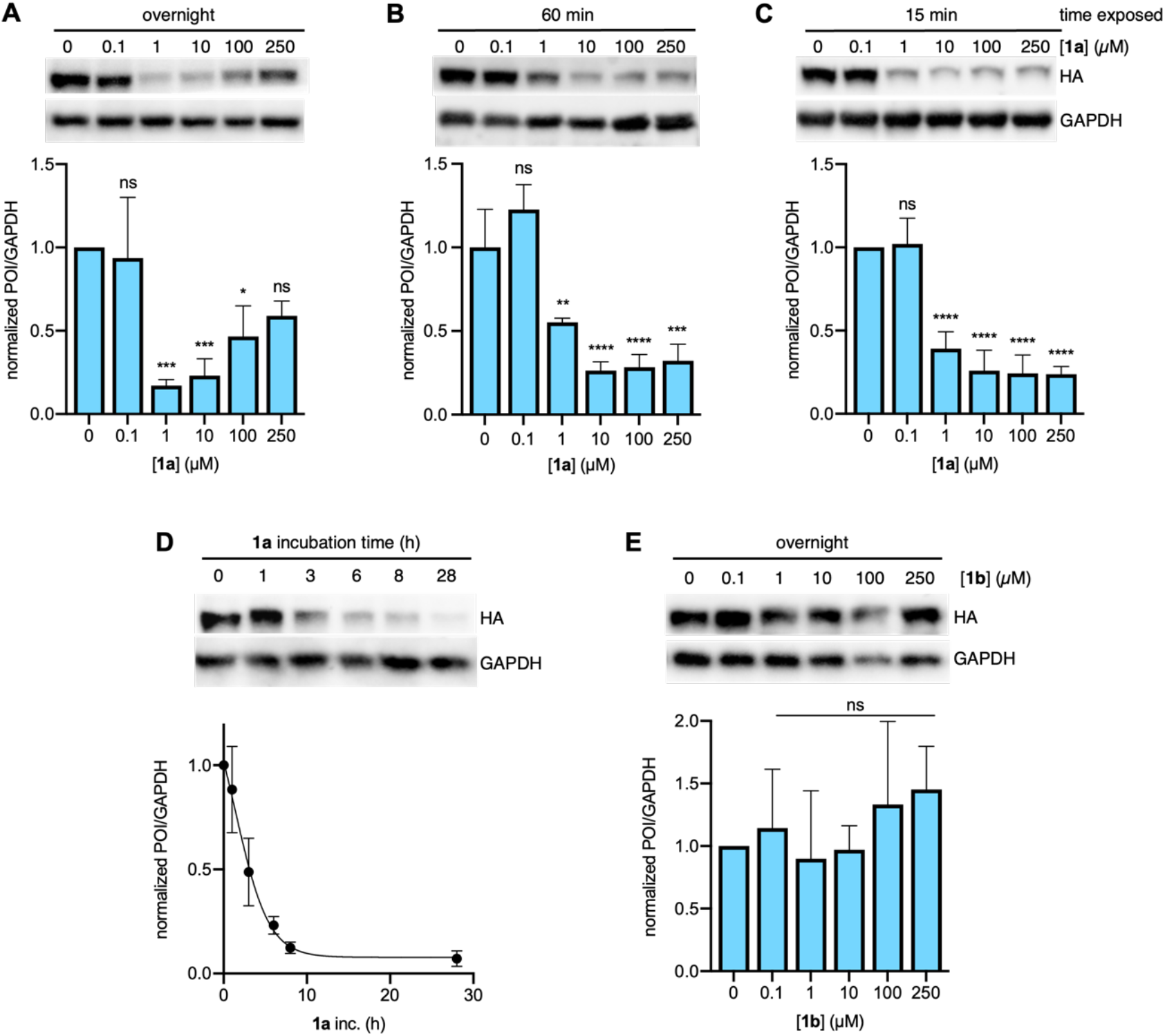
Optimization of bioorthogonal E3 ligase mediated degradation in mammalian cells. Effect of increasing concentrations of **1a** on mCherry-EGFP-Y151TetF-HA degradation after A) overnight, B) one hour, or C) 15 min exposure to the compound, each incubated overnight. D) Time course of **1a** (10 µM) mediated degradation of mCherry-EGFP-Y151TetF, as determined by two biological replicates analyzed by western blot. E) Comparison of the effect of PEG linker length on degradation via a dose response of mCherry-EGFP-Y151TetF treated with increasing concentrations of **1b**. A-C, and E were performed in triplicate with error bars representing standard deviation from the mean. Statistical significance was determined by one-way ANOVA relative to the negative control, where ^*^ = p < 0.1, ^**^ = p < 0.01, ^***^ = p < 0.001, and ^****^ = p < 0.0001. Full blots used for quantification are available in **Fig. S4**.

In an attempt to address the observed hook effect, we removed excess **1a** from cells after protein conjugation. Specifically, the transfected cells were compound-treated for the indicated time prior to placement into CRBN ligand-free media for subsequent overnight incubation. By reducing the total time exposed to **1a** from overnight to just 15 or 60 minutes, we observed a distinct reduction of the hook effect (Fig. 2B-C). While degradation was still optimal following overnight treatment with 1 or 10 µM of **1a**, the significant target degradation despite dramatically decreased ligand exposure times indicates that the bioconjugation reaction was fast, leaving the UPS-mediated degradation as the limiting factor. The kinetics of the degradation were further validated by a time-course, in which cells expressing the TetF reporter were incubated with 10 µM of **1a** for the indicated times (Fig. 2D). Substantial degradation was observed within just 6 hours, with complete knockdown achieved after overnight incubation. To account for variations in levels of TetF-bearing proteins expressed in cells, subsequent degradation experiments were performed with overnight exposure to 10 µM of **1a**. Finally, the specificity of **1a** for TetF-bearing proteins was confirmed, as no degradation of wild-type mCherry-EGFP was observed after overnight exposure to **1a** (Fig. S3).

Following successful degradation with **1a**, we wanted to compare the degradation efficiency of a bioorthogonal E3 ligase ligand bearing a markedly longer PEG-linker, as linker length is known to affect PROTAC potency and efficiency.^59,55,54,56^ Thus, we synthesized sTCO-PEG9-CRBN (**1b**) and found that it surprisingly did not induce any significant degradation of the mCherry-EGFP-Y151TetF target (Fig. 2E). A pulse-chase experiment with **1b** validated 85% bioconjugation, so degradation was not limited by the covalent attachment of **1b** to the protein surface (Fig. 3D). We instead hypothesized that this observed lack of activity is due to the spatial orientation of the resultant ternary complex, as the PEG9 linker is likely too long to induce a productive E3 ligase complex of this small protein target. While **1b** did not exhibit degradation of mCherry-EGFP-Y151TetF, we hypothesized that the longer linker might yield improved degradation of different sites on the same reporter or for other larger, or more complex protein targets, so we moved forward with both compounds.

**Fig 3.**
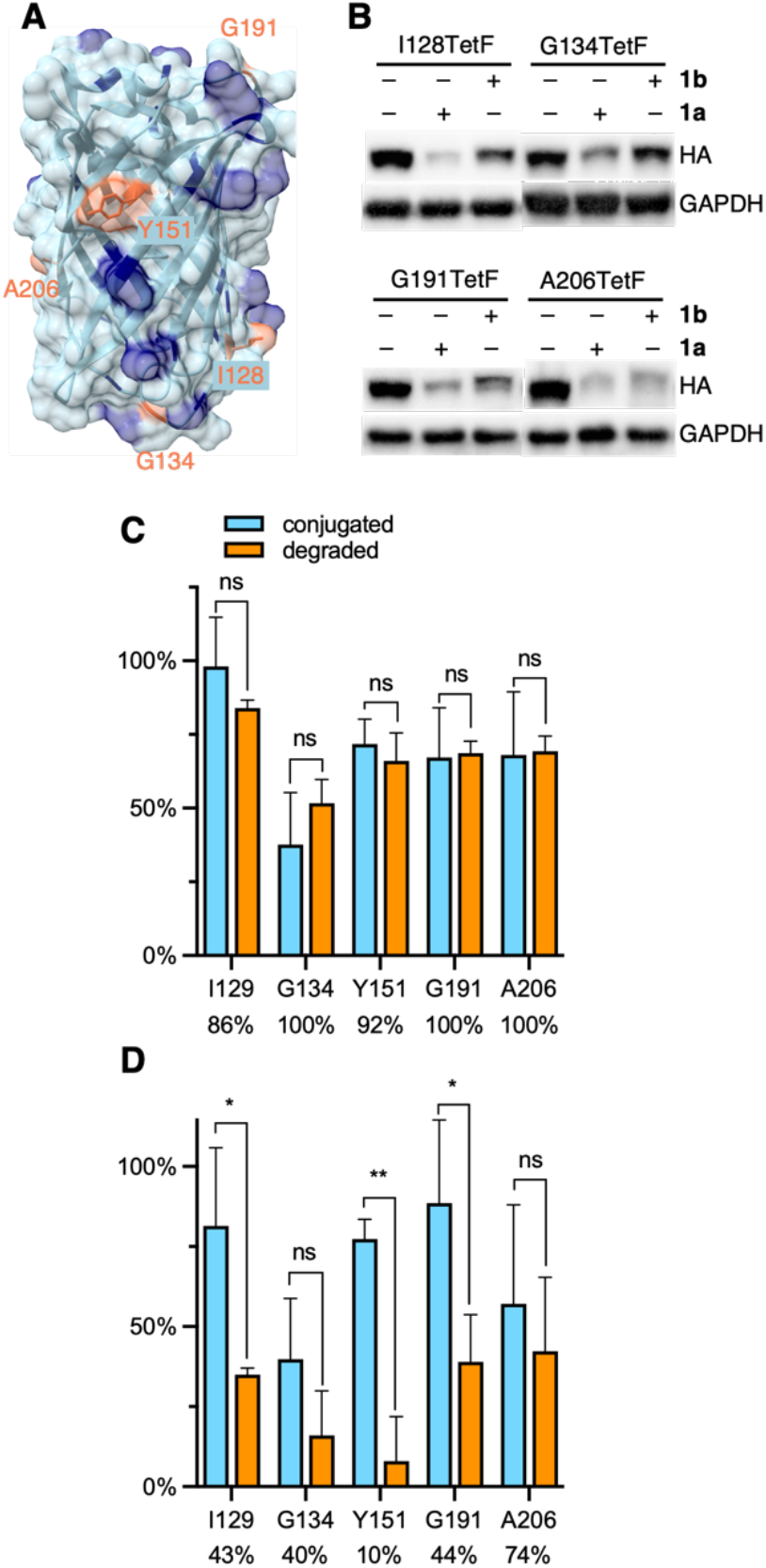
A) Crystal structure of EGFP with mutated residues highlighted in orange and lysine residues highlighted in blue (PDB 2Y0G). B) Representative western blots of mCherry-EGFP-TetF mutants following treatment with **1a** or **1b**. Full blots are available in Fig. S10. Quantification of C) **1a** and D) **1b** conjugation versus degradation observed for each EGFP-TetF mutant, with percentage of degradation per bioconjugated protein listed below each mutant. Percent conjugation was determined by pulse-chase analysis in the presence of proteasome inhibitor MG132. Error bars were calculated as standard deviation from the mean of three biological replicates with statistical significance determined via unpaired t-test in Prism 10, where ^*^ = p < 0.1 and ^**^ = p < 0.01.

A distinct advantage of our E3 ligase ligand conjugation approach compared to small molecule PROTACs or dTAGs is the ability to directly compare degradation of ligase recruitment to different locations across a protein surface. This feature is particularly useful for the analysis of proteins without known small molecule ligands, as a protein surface region that yields successful degradation can be specifically targeted for small molecule ligand discovery. EGFP has no known small molecule ligand. Thus, with our system optimized to degrade mCherry-EGFP-Y151TetF-HA, we designed four additional EGFP mutants, I128TetF, G134TetF, G191TetF, and A206TetF (**Fig. 3A**), to determine if different sites of ligand recruitment led to varying levels of degradation.

Before comparing the degradation of different mCherry-EGFP mutants by **1**, we first needed to validate the extent of bioconjugation to TetF at each position. To accurately determine the bioconjugation efficiency of **1a** and **1b**, we developed a pulse-chase assay. Cells expressing the TetF-bearing protein are “pulsed” with **1a** or **1b** (10 µM) for 1 hour, then “chased” with sTCO-CA (10 M) for 1 hour. Proteins not conjugated by **1** are available for reaction with sTCO-CA, for subsequent covalent heterodimer formation with HaloTag following cell lysis. Western blot analysis is then used to determine the extent of conjugation to sTCO-CA (**Fig. S1B**), which reveals the amount of protein bioconjugated by **1**. Due to the potential for degradation by **1** within the 2 hour timeframe of the experiment, these pulse-chase analyses are performed in the presence of the proteasome inhibitor MG132.

Using this pulse-chase method, we identified respective labeling efficiencies of **1a** and **1b** for TetF-mutants of mCherry-EGFP-HA. The observed bioconjugation of **1a** and **1b**, respectively, were 98% and 81% (I129TetF, **Fig. S5**), 38% and 40% (G134TetF, **Fig. S6**), 72% and 77% (Y151TetF, **Fig. S7**), 67% and 89% (G191TetF, **Fig. S8**), and 68% and 57% (A206TetF, **Fig. S9**).

By determining the extent of bioconjugation to TetF mutants across the surface of EGFP, we were able to then account for variations in overall degradation relative to the amount of **1** conjugated to the POI (**Fig. S10**). For all five mCherry-EGFP-TetF-HA mutants examined, we observed that nearly every protein that was reacted with **1a** was degraded (**Fig. 3C**). Interestingly, poor degradation was observed with **1b** for I129TetF, G134TetF, Y151TetF, and G191TetF, but moderate degradation was afforded for A206TetF (**Fig. 3D**).

The small size and abundance of surface lysine residues made EGFP an ideal target for initial methodology validation. To expand the scope of protein degradation through E3 ligase ligand conjugation, we selected another small protein without a known PROTAC, the mitogen-activated protein kinase dual specificity phosphatase 6 (DUSP6, Fig. 4A). DUSP6 is involved in the Ras/Raf/MAPK pathway, in which it dephosphorylates MAP kinases ERK, p38, and JNK.^60,61^ Due to the high degree of conservation between structures of DUSP6 and other dual specificity phosphatases, the discovery of specific small molecule ligands has been limited.^62^ For example, DUSP6 has one known allosteric inhibitor, BCI, however this ligand also recognizes and inhibits DUSP1.^62,63^ Because DUSP6 plays critical roles in signaling processes that affect development, differentiation, and proliferation, we selected it as another model for probing a previously undruggable and “un-PROTAC-able” protein.

**Fig 4.**
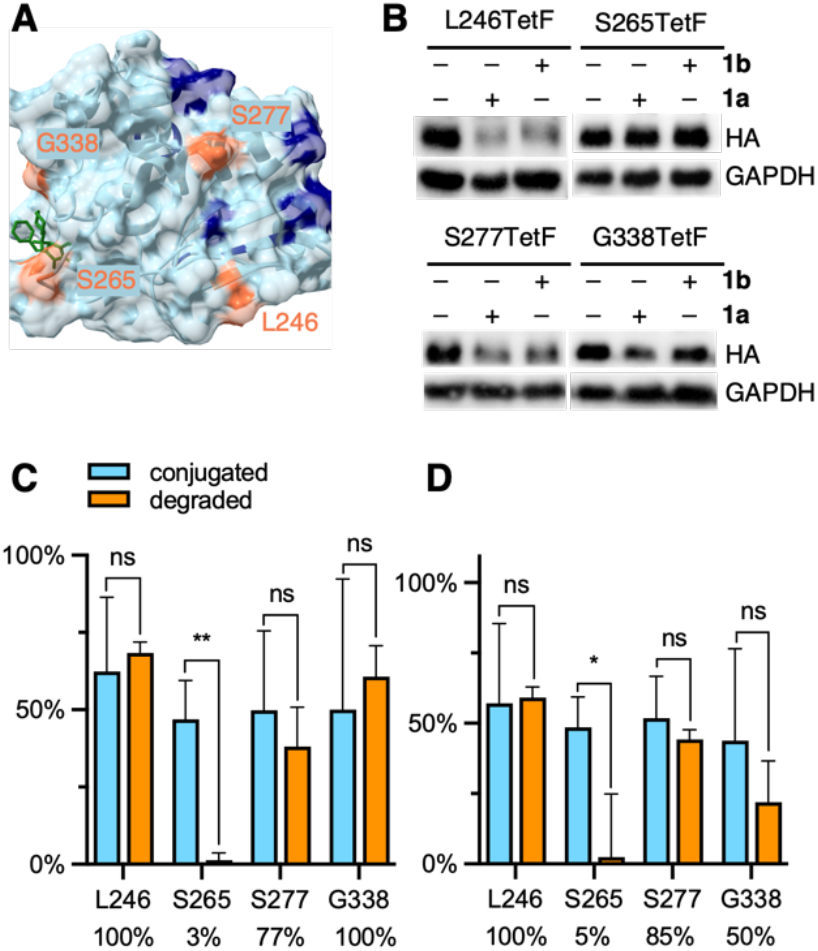
A) Crystal structure of DUSP6 with mutated residues highlighted in orange and lysine residues highlighted in blue (PDB 1MKP), docked with the allosteric ligand BCI (green structure). B) Representative western blots of DUSP6-TetF mutants following treatment with **1a** or **1b**. Full blots are available in Fig. S15. Quantification of C) **1a** and D) **1b** conjugation versus degradation observed for each DUSP6-TetF mutant, with percentage of degradation per bioconjugated protein listed below each mutant. Percent conjugation was determined by pulse-chase analysis in the presence of proteasome inhibitor MG132. Error bars were calculated as standard deviation from the mean of three biological replicates with statistical significance determined via unpaired t-test in Prism 10, where ^*^ = p < 0.1 and ^**^ = p < 0.01.

We generated four TetF surface mutants based on the crystal structure of DUSP6. We selected two mutants, L246TetF and S277TetF, for their close proximity to lysine residues and relative positioning away from the BCI binding site. We further selected two mutants near the BCI binding site, S265TetF and G338TetF (Fig. 4A). We then confirmed and quantified bioconjugation to each DUSP6-TetF mutant utilizing the abovementioned pulse-chase analysis, pulsing with either **1a** or **1b**, and chasing with sTCO-CA. The observed bioconjugation of **1a** and **1b**, respectively, were 62% and 57% (L246TetF, Fig. S11), 47% and 49% (S265TetF, Fig. S12), 50% and 52% (S277TetF, Fig. S13), and 50% and 44% (G338TetF, Fig. S14).

Utilizing the previously optimized conditions for degradation through bioconjugation of **1**, we found that the DUSP6-L246TetF mutant was completely degraded by both **1a** and **1b** (Fig. 4B-D). As the incorporated tetrazine is situated near (within 10-20 angstroms) 2-3 surface lysine residues, the observed degradation aligns with the known mechanism of ubiquitination.^64^ Interestingly, the other mutation located in a lysine-dense region, S277TetF, showed 77% or 85% degradation relative to the extent of **1a** or **1b** conjugation, respectively (Fig. 4C-D). This incomplete degradation upon CRBN recruitment to S277TetF-DUSP6 indicates that lysine density and proximity may not be the sole predictors of an excellent PROTAC, further validating the need for this methodology.

With evidence that DUSP6 can be degraded by CRBN, we next looked to the mutants positioned near the BCI binding domain to determine the PROTAC-ability of the ligand. The S265TetF-DUSP6 mutant yielded moderate bioconjugation by **1a** and **1b** (Fig. 4C and D, respectively) but no degradation was observed after exposure to either compound. Based on the distance (> 20 angstroms) of the S265 residue from accessible surface lysines, the lack of UPS-mediated degradation is unsurprising. The G338 residue is near surface K324, which is likely the reason for the observed degradation by **1a** (Fig. 4C). Interestingly, despite no degradation observed with longer PEG lengths in initial EGFP studies, the amount of DUSP6 degradation relative to bioconjugation of **1b** was comparable to **1a** for S277TetF-and L246TetF-DUSP6, with G338TetF-DUSP6 degradation efficiency decreasing. Despite the trend in decreased degradation for longer linkers (>8 atoms) of CRBN-recruiting PROTACs,^65^ this observed trend between **1a** and **1b** suggests some flexibility in DUSP6 PROTAC design. Further, these trends support the need for a streamlined approach to PROTAC-component screening. Finally, based on the poor degradation through recruitment of CRBN to DUSP6 mutant S265TetF, we suspect that BCI, even with improved specificity for DUSP6, would not yield a potent CRBN-based PROTAC. Instead, a PROTAC binder to the L246/S277 region of the protein should be developed, which would first require discovery of a suitable small molecule ligand.

In addition to protein targets without known ligands, we sought to test our bioorthogonal E3 ligase ligand on a target with a known PROTAC. The estrogen related receptor alpha (ERRα) is an orphan nuclear hormone receptor and its PROTAC consists of a diarylether thiazolidenedione ligand (Fig. 5A, green structure) linked via a short hydrocarbon to the peptidomimetic ligand that recruits the VHL E3 ligase.^6^ The ERRα ligand of the PROTAC undergoes a reversible covalent reaction with ERRα^66^ that leads to high selectivity and potent degradation (at 1 µM).^6^ PROTACs bearing the same ERRα ligand linked to ligands for CRBN, MDM2, and IAP yielded no target degradation.^67^ Therefore, we aimed to investigate if ERRα could be degraded by an E3 ligase ligand other than VHL if it was recruited to a completely different region of the protein surface. To test this, we first generated four mutants of ERRα for the genetic incorporation of TetF on the protein surface. Two mutations, F382TetF and F495TetF, were selected for their positions in the known ERRα PROTAC binding domain (Fig. 5A). Two additional mutants, V299TetF and H437TetF, were selected because of their distance away from the known PROTAC binding location. We validated bioconjugation efficiency of each mutant using the same pulse-chase method described above, before moving to degradation studies. The observed bioconjugation yields of **1a** and **1b**, respectively, were 36% and 31% (V299TetF, Fig. S16), 37% and 28% (F382TetF, Fig. S17), 32% and 29% (H437TetF, Fig. S18), and 30% and 43% (F495TetF, Fig. S19).

**Fig 5.**
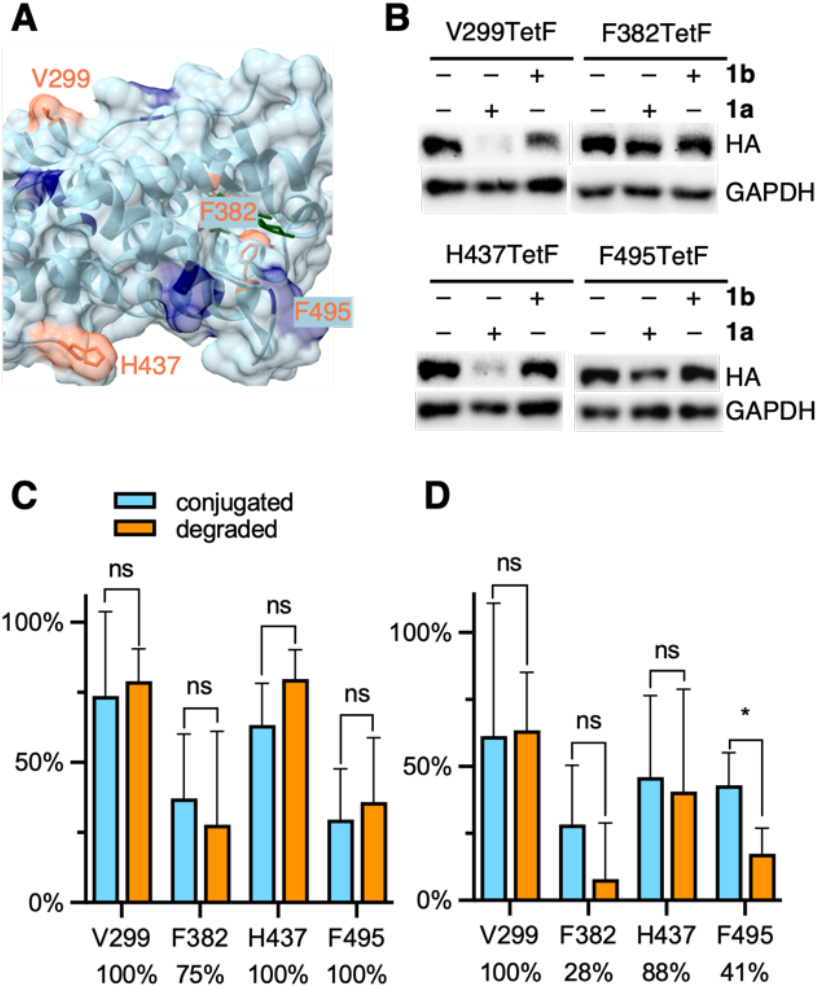
A) Crystal structure of ERRα with mutated residues highlighted in orange, lysines highlighted in blue, co-crystalized with the known ERRα ligand in dark green (PDB 3K6P). B) Representative western blots of ERRα-TetF mutants following treatment with **1a** or **1b**. Full blots are available in Fig. S19. Quantification of C) **1a** and D) **1b** conjugation versus degradation observed for each ERRα-TetF mutant, with percentage of degradation per bioconjugated protein listed below each mutant. Percent conjugation was determined by pulse-chase analysis in the presence of proteasome inhibitor MG132. Error bars were calculated as standard deviation from the mean of three biological replicates with statistical significance determined via unpaired t-test in Prism 10, where ^*^ = p < 0.1 and ^**^ = p < 0.01.

Despite relatively low amounts of **1a** or **1b** bioconjugation observed for each mutant of ERRα, the amount of protein degraded relative to the amount of observed conjugation revealed interesting trends. F382TetF-and F495TetF-ERRα, the two PROTAC binding pocket mutants, demonstrated moderate to excellent degradation efficiency for **1a**, but poor degradation for **1b** (Fig. 5C-D). Previous SAR studies observed no degradation of ERRα with CRBN-, MDM2-, or IAP-based ERRα-PROTACs but successful degradation via VHL recruitment.^67^ Our demonstration of CRBN-mediated degradation through bioorthogonal labeling of the PROTAC binding sites with a cereblon-recruiting ligand indicates potential for expanding the chemistry of these PROTACs, further validating the need for rapid, straightforward comparison of a target’s “PROTAC-ability” using the currently available systems.

The two mutants bearing surface residues not associated with the VHL-based PROTAC binding region, V299TetF-ERRα and H437TetF-ERRα, were completely degraded by **1a**. Further, complete degradation was observed for V299TetF-ERRα bioconjugated with **1b** (Fig. 5B and D), whereas the amount of **1b**-mediated degradation decreased for H437TetF-ERRα (88%, Fig. 5B and D). These observed variations in CRBN-mediated degradation at different locations on ERRα indicate a previously uncharacterized correlation between E3 ligase recruitment location and degradability by ligand type. Thus, POIs previously not degraded by thalidomide-based PROTACs could potentially be degraded by simply changing the location of target binding, further expanding the scope of future PROTACs for ERRα.

## Conclusion

In conclusion, we have developed a conjugatable E3 ligase ligand for the efficient degradation of multiple proteins of interest (POIs), including the fluorescent reporter EGFP, the phosphatase DUSP6, and the hormone receptor ERRα. These proteins were site-specifically modified with a reactive bioconjugation handle for rapid bioorthogonal modification with said E3 ligase ligand in human cells. The optimal conditions for protein degradation after conjugation with sTCO-PEGn-CRBN (**1**) were determined to be overnight, as degradation was limited by the capacity of the proteasomal machinery, not the fast bioconjugation reaction. Additionally, incomplete bioconjugation, which was quantified through pulse-chase analysis for each mutant of each protein, impacted the observed degradation in several cases, albeit ∼75-90% bioconjugation yields were achieved. Our investigations also showed differential degradation for mutants of three different proteins when shorter (PEG2 vs PEG9) linkers were used between the conjugation handle and the E3 ligase ligand. As linker length and composition are known to affect PROTAC specificity and potency,^55,56,68^ our method for screening linker length independent of POI binding will streamline future PROTAC designs.

Four TetF-bearing mutants of EGFP were degraded by **1a**, demonstrating the versatility of this tool for inducing degradation through attachment at different regions of a protein surface and for a protein without a known small molecule ligand. The recruitment of the E3 ligase cereblon was further demonstrated for TetF-mutants of the dual specificity phosphatase DUSP6 and the hormone receptor ERRα. DUSP6, which has no known PROTAC, was not degraded when the E3 ligase ligand was recruited to a known small molecule binding location, but instead showed excellent degradation when recruited to a lysine-rich region of the protein surface. This discovery validates that DUSP6 can be degraded, but that its known small molecule inhibitor will be a poor ligand to initiate PROTAC design. Our approach instead highlighted new hotspots for DUSP6 degradation, which we believe will assist in guided ligand design. ERRα bioconjugation showed limited yields, but excellent degradation was observed for proteins bioconjugated to **1a**. These findings indicate the potential for development of a CRBN-recruiting ERRα PROTAC in addition to the previously reported VHL-PROTAC. Expansion of the ERRα PROTAC toolbox may be possible through recruitment of cereblon to different regions of the protein surface, as indicated by the excellent degradation of ERRα mutants V299TetF and H437TetF.

The targeted E3 ligase ligand positioning on any POI opens several avenues for PROTAC development and expansion, such as the capitalization on tissue-specific E3 ligases and optimization of the stability and pharmacokinetics of PROTACs.^69,70^ Using the flexibility and complete site-specificity offered by genetic code expansion for probing different protein surface locations we hope to guide novel ligand design by prioritizing regions within the protein of interest for the discovery of small molecule binders. Further, by directly comparing degradation at different sites with a collection of linkers and E3 ligase ligands, we provide a new tool to expand the “PROTAC-ability” of new targets and improve upon previously “unPROTACable” proteins.

In addition to the ability to investigate proteins and distinct sites on their surfaces for PROTAC design, our bioorthogonal E3 ligase ligand can be used as a generalizable OFF switch for protein function. Our method bypasses the need for protein-specific small molecules required for classical PROTAC approaches, lessening the required synthetic workload and streamlining the study of novel and promiscuous proteins. Further, we are able to knock down proteins without the need for protein fusions to recruit heterobifunctional degraders, as is required by systems like dTAGs^8,9^ or the peptide-based HiBiT-SpyTag system.^71^ Paired with the ability to temporally control POI expression through **TetF** additional, a conditional ON/OFF switch system was developed. The use of genetic code expansion paired with a small, site-specific bioorthogonal handle provides no or minimal perturbation of native protein function.

With the validation of this method for probing the PROTACability of any POI, we aim to expand our toolbox of bioorthogonal E3 ligase ligands to include ligands for other E3 ligases like VHL, MDM2, and IAP, as well as different linker chemistries, and compositions for facile elucidation of optimal PROTAC design for new protein targets. This broad applicability of the approach is supported by a recent site-specific installation of hydrophobic ligands on a protein of interest for targeted degradation, that was reported over the course of our studies.^72^ As the field of targeted protein degradation continues its rapid growth, our approach for the bioorthogonal recruitment of E3 ligase ligands to any protein of interest will streamline PROTACability studies and advance PROTAC applications.

## Author Contributions

Conceptualization: A. D.; methodology: O. S., G. B., A. R.; writing: O. S. and A. D.

## Data Availability

The biological and chemical experimental protocols, supplementary figures and tables, and NMR spectra are available in the supplementary information.

## Acknowledgements

This work was supported by the National Science Foundation (CHE-1904972) and aided by GCE4All Biomedical Technology Development and Dissemination Center supported by the National Institute of General Medical Science (RM1GM144227), specifically by supplying the synthetase plasmid pAcBac-TetFRS. We would like to thank Dr. Mark Howarth for depositing Addgene plasmid 105627 (HaloTag), Dr. Toren Finkel for depositing Addgene plasmid 10975 (ERRα), and Dr. Dustin Maly for depositing Addgene plasmid 40772 (DUSP6). Some figure elements were created with BioRender.com.

