## Supplemental Information for "Investigating Protein Degradability through Site-Specific Ubiquitin Ligase Recruitment"

### 1. Biological Protocols

#### 1-1a. Plasmid Construction: General Protocol

All site-directed mutagenesis reactions were performed using the same protocol with varying primers, template DNA, annealing temperatures, and elongation times. Mutagenesis was completed in a 50 µL reaction containing template DNA (10 ng), 10 µL of 5x NEB GC buffer (Thermo Scientific, F530L), 1 µL of dNTP mix (10 mM, Thermo Scientific, R0191), 1 µL of both forward and reverse primers (5 µM, Sigma Aldrich, see sequence in **Table S1**), 0.75 µL of DMSO, and 0.5 µL of Phusion polymerase (2 U/µL, Thermo Scientific, F530L). Reactions were performed using an initial denaturing step (95 °C, 3 minutes), followed by 30 cycles of denaturing (95 °C, 30 seconds), annealing (see temperatures in **Table S1**), and elongation (72 °C, see times in **Table S1**) steps. The thermocycler program was completed with a final elongation step (72 °C, 10 minutes), gradually cooled (16 °C, 5 minutes), and the samples were stored at –20 °C until further use. Template DNA was digested by adding 5 µL 10x CutSmart buffer and 1 µL of DpnI (20 U/µL, NEB, R0176S) to the 50 µL PCR mixture, and incubating for 1 hour at 37 °C.

To assess the purity of the PCR product, 2 µL of the mixture was diluted in 8 µL of water and 2 µL of 6x loading dye (NEB, B7024S) and run on a 0.8% (w/v) agarose gel in Tris-buffered EDTA (TBE, 10 mM Tris-HCl, 1 mM EDTA, pH = 8.0) buffer at 80 V for 30 minutes. If a single band corresponding to the correct length was observed, the mixture was used immediately for transformation. If impurities were observed, the PCR product was purified by gel electrophoreses of the rest of the sample, band excision, and gel extraction.

To gel extract, 10 µL of 6x loading dye was added to the DpnI-treated PCR mixture, and the entire amount was run on a 0.8% (w/v) agarose gel. The band corresponding to the correct length was cut from the gel and purified with the GeneJET Gel Extraction Kit (Thermo Scientific, K0692) according to the manufacturer's protocol.

To transform cells, a 50 µL aliquot of Top10 competent cells was thawed on ice in a 1.7 mL microcentrifuge tube (Laboratory Products Sales, L211511) and combined with 5 µL of the crude PCR mixture, or 75 ng of purified plasmid. The mixture was incubated on ice for 30 minutes. The cells were heat shocked by incubating the tube in a hot water bath (42 °C, 30 s), followed immediately by incubation on ice for 2 minutes. Ice-cold SOC media (200 µL, VWR, 100219-988) was applied to the cell mixture, and the tube was incubated at 37 °C with shaking at 250 rpm for 1 hour. The entire contents of the transformation mixture were plated on 10 mL of LB agar supplemented with 10 µL of a 100 mg/mL antibiotic (ampicillin or kanamycin) stock and incubated overnight at 37 °C. Three colonies were inoculated into 5 mL of LB broth supplemented with 5 µL of a 100 mg/mL antibiotic (ampicillin or kanamycin) stock and grown overnight with shaking (37 °C, 250 rpm), followed by DNA miniprep (Thermo Scientific, K0503). Plasmid sequences were confirmed by Genewiz Sanger sequencing using universal CMV forward and EGFP-C reverse primers (P1 and P2, respectively, **Table S1**).

All Gibson assembly cloning procedures were performed using the following protocol with varying template DNA, annealing temperatures, and elongation times.[1] Backbone and insert amplifications were completed in a 100 µL reaction containing template DNA (20 ng), 20 µL of 5x NEB HF buffer, 2 µL of dNTP mix (10 mM, each, Thermo Scientific, R0191), 1 µL each of forward and reverse primers (10 µM, each, Sigma Aldrich, sequences, annealing temperatures, and elongation times available in **Table S1**), 1.5 µL of DMSO, and 1 µL of Phusion polymerase (2 U/µL, Thermo Scientific, F530L). The thermocycler program outlined above was used for insert and backbone amplification, and the products were gel extracted as described above.

The DNA fragments were annealed using the published Gibson assembly method.[1] The 5x isothermal reaction buffer was prepared (25% PEG-8000 (Alfa Aesar), 500 mM Tris-HCl (Amresco) pH 7.5, 50 mM $MgCl_2$ (Alfa Aesar), 50 mM DTT (Fisher), 1 mM each of the 4 dNTPs (Fisher), and 5 mM NAD (NEB)) and stored at –20 °C. The master mix was prepared (86 µL of autoclaved MilliQ water, 36 µL of 5x isothermal reaction buffer, 2 µL of Phusion DNA polymerase (2 U/µL Fisher), 0.064 µL of T5 exonuclease (10 U/µL, NEB), and 0.016 µL of Taq ligase (40 U/µL, abm)) and stored at –20 °C in 15 µL aliquots before use. Final DNA products were transformed, purified, and sequenced as described above.

**1-1b. Plasmid Construction: pmCherry-EGFP-HA-TAG**

Using the Gibson assembly protocol outlined in the Plasmid Construction: General Protocol section, the mCherry gene was amplified from pmCherry-N1 (Clontech, 632523) using primers P3 and P4 and cloned into the pEGFP-HA backbone (amplified with P5 and P6) for assembly of pmCherry-EGFP-HA. The site-directed mutagenesis protocol described in section **1.1a** was used to introduce the following mutations: Y151TAG (P7 and P8), I128TAG (P9 and P10), G134TAG (P11 and P12), G191TAG (P13 and P14), A206TAG (P15 and P16).

**1-1c. Plasmid Construction: pDUSP6-HA-TAG**

EGFP was removed from pWT-DUSP6-EGFP-HA[2,3] using primers P17 and P18 to yield pWT-DUSP6-HA. The site-directed mutagenesis protocol described in **1-1a** was used to introduce the following mutations into pDUSP6-HA: L246TAG (P19 and P20), S265TAG (P21 and P22), S277TAG (P23 and P24), G338TAG (P25 and P26).

**1-1d. Plasmid Construction: pERRα-HA-TAG**

Using the Gibson assembly protocol outlined in the Plasmid Construction: General Protocol section, the ERRα gene was amplified from pCMV-flag-ERRalpha[4] (Addgene, 10975) using primers P27 and P28 and cloned into the pEGFP-HA backbone (P29 and P30) for assembly of pERRα-EGFP-HA. The EGFP gene was removed through using primers P31 and P32 for generation of pERRα-HA. The site-directed mutagenesis protocol described in section **1.1a** was used to introduce the following mutations: V299TAG (P33 and P34), F382TAG (P35 and P36), H437TAG (P37 and P38), F495TAG (P39 and P40).

#### 1-1d. Plasmid Construction: pE323-TetFRS

The TetFRS insert was amplified from pAcBac-TetFRS[5] using primers P41 and P42 using the PCR protocol outlined in 1-1a Plasmid Construction: General Protocol. The pE323 backbone was prepared by first diluting 10000 ng of pE323-HCKRS[6] with water to a final volume of 43 µL. The plasmid was further diluted with 5 µL of CutSmart buffer, then digested with 1 µL of XbaI (NEB, R0145S) and 1 µL of NotI-HF (NEB, R3189L) restriction enzymes. The digest was done simultaneously with alkaline phosphatase treatment by adding 5 µL of AP reaction buffer (NEB, M0289S) and 1 µL of Antarctic phosphatase (NEB, M0289S) directly to the mixture. The reaction was incubated at 37 °C for 2 hours.

The PCR amplified insert and the restriction digested backbone were both gel extracted from an agarose gel, as described above. The purified insert was ligated into the purified backbone by incubating 430 ng of the backbone with 660 ng of the insert with 1 µL of T4DNA ligase buffer (NEB 50811605) and 0.5 µL of T4 DNA ligase (NEB 50811605). The reaction was incubated overnight at 4 °C. The success of the ligation was confirmed via agarose gel then the DNA was transformed into Top10 competent cells as described above. pE323-TetFRS and pAcBac-TetFRS displayed the same levels of transfection efficiency in mammalian cells and were thus used interchangeably.

### 1-2. Cell Culture

All cell culture experiments were performed in a sterile laminar flow hood. HEK293T cells (ATCC, CRL-11268) were maintained in Dulbecco's Modified Eagle Medium (DMEM, Gibco, SH30003.03) supplemented with 10% (v/v) fetal bovine serum (Sigma Aldrich, F0926) and 1% (v/v) penicillin/streptomycin (complete medium, Corning, 30002CI) at 37 °C with 5% $CO_2$. Cells were used between passage number 3 and 32 and tested for mycoplasma contamination every 6 months (Genlantis, MY01100). All cell culture medias are warmed to 37 °C prior to application to cells.

### 1-3. Western Blot

#### 1-3-1. Transfection

HEK293T cells were seeded into a poly-D-lysine (PDK) treated 48-well plate (50,000 cells/well, Greiner, 677180) in 250 µL of complete medium. When cells reached 80-90% confluency, the media in each well was replaced with 200 µL of fresh, antibiotic-free DMEM supplemented with 0.25 mM tetrazine phenylalanine (**TetF**[7]). The UAA containing media was made by diluting 12.5 µL of a 100 mM stock solution into 5 mL of antibiotic-free DMEM to create a master solution containing a final concentration of 0.25% DMSO.

A master mix transfection solution was prepared by combining 400 ng of DNA per well (1:2 TAG-mutant/TetFRS) in 40 µL of OptiMEM transfection media (Thermo Scientific, 22600050) and 2 µL of 1 mg/mL linear polyethyleneimine (LPEI, Polysciences, 23966) per well. For example, transfection of six wells in a 48-well plate requires 800 ng of TAG-containing plasmid (pmCherry-EGFP-TAG-HA, pDUSP6-HA-TAG, or pERRα-HA-TAG) and 1600 ng of pAcBac-TetFRS diluted into 240 µL of OptiMEM and 12 µL of LPEI. The transfection master mix was gently mixed by inversion and incubated for 10 minutes at room temperature. Following incubation, 40 µL of the mixture was added dropwise to each well. The cells were then incubated at 37 °C and 5% $CO_2$. for 24 hours.

Following overnight transfection, the cells were washed by gently applying 200 µL of fresh complete medium to each well and immediately removing the media. A second wash was completed by adding another 200 µL of fresh complete medium to each well and incubating the cells at 37 °C for 30 minutes, and gently removing the supernatant.

#### 1-3-2. Compound Treatment

Quantification of bioconjugation to TetF-bearing proteins was performed similarly to the previously described method, adapted to a pulse-chase format.[8] Following transfection with pE323-TetFRS and the plasmid encoding the TAG-containing protein of interest, cells were washed once with 200 µL of fresh complete medium. To limit any degradation caused by **1**, the initial wash solution was removed, and cells were then treated with 200 µL of fresh medium supplemented with 10 µM of the proteasome inhibitor MG132 and incubated at 37 °C for 4 hours. Next, a working solution of the 'pulse' treatment was prepared by diluting 4 µL of 10 mM of MG132 and 4 µL of 10 mM of **1a**, **1b**, or DMSO into 4 mL of fresh complete medium. The cells wer[1]e then treated with 200 µL of the pulse solution for 1 hour at 37 °C. The chase solution was then prepared by diluting 4 µL of 10 mM of MG132 and 4 µL of 10 mM of sTCO-CA or DMSO into 4 mL of fresh complete medium. The cells were treated with 200 µL of the chase solution for 1 h at 37 °C. Finally, cells were washed with 200 µL of cold PBS (5 min) and then lysed (**1-3-3**). The cell lysate (60 µL) was combined with 5 µL of an 80 µM solution of HaloTag[8,9] and incubated at 37 °C overnight.

For **sTCO-PEG$_n$-CRBN** (**1**) dose-response experiments (**Figure 2A**), cells were transfected as described above and a 250 µM working solution of **1** was made by diluting 25 µL of a 10 mM stock (in DMSO) into 975 µL of complete medium. To make a 100 µM mixture, 400 µL of the 250 µM stock was diluted into 600 µL of complete medium. Further 10x serial dilutions were made by diluting 100 µL of the 100 µM solution into 900 µL of complete medium. This serial dilution was repeated to generate the designated concentrations. Following the removal of TetF after transfection, 200 µL of media supplemented with or without **1** was applied to each well and the plate was incubated at 37 °C with 5% $CO_2$ overnight. Cells were then lysed (**1-3-3**) and analyzed by western blot (**1-3-4**).

For mechanistic insight into **1a** degradation (**Fig. S2A**) cells were transfected as described above. A 10 µM working solution of media containing either MG132 or epoxomicin was made by adding 1 µL of the respective proteasome inhibitor (10 mM in DMSO) into 999 µL of complete medium. Following removal of TetF, 200 µL of media supplemented with either MG132, epoxomicin, or DMSO was added to each well and the cells were incubated for 4 h. Next, a solution containing 50 µM of 1a was made by adding 5 µL of **1a** (10 mM in DMSO) into 995 µL of complete medium.

Following the allotted time for proteasome inhibitor incubation, 50 µL of 50 µM **1a** was added directly to each well, for a final volume of 250 µL and final concentration of 10 µM of **1a**. The cells were then incubated overnight at 37 °C with 5% $CO_2$ before lysis (**1-3-3**) and western blot analysis (**1-3-4**).

Tetrazine specificity of **1a** (**Fig. S2B**) was confirmed following transfection of HEK293T cells with wild-type pmCherry-EGFP-HA (no TAG mutant). Transfection was performed similarly to the method described above. Instead, no TetF was included in the antibiotic-free DMEM, only 133 ng of pmCherry-EGFP-HA was added per well, and no additional synthetase plasmid was included in the transfection mastermix. Following overnight expression, cells were similarly washed before incubation with 10 µM of **1a** overnight. Cells were lysed (**1-3-3**) and resultant protein levels were analyzed by western blot (**1-3-4**).

For **1a** hook effect mitigation (**Figure 2B-C**), the same concentrations of **1a** were made using the same dilutions as above. For each condition, 200 µL of supplemented media was added to the respective well and incubated for either 15 or 60 minutes. The media containing **1a** was removed, and 200 µL of fresh complete medium was applied. The cells were then incubated overnight at 37 °C with 5% $CO_2$ before lysis (**1-3-3**) and western blot analysis (**1-3-4**).

For degradation of different TetF-bearing proteins by **1**, a 10 µM solution of **1** was made by diluting 3 µL of **1** (10 mM in DMSO) into 3 mL of pre-warmed complete medium. Following TetF washout, 200 µL of medium supplemented with or without 1 was added to each well and the plate was incubated at 37 °C and 5% $CO_2$ overnight before lysis (**1-3-3**) and western blot analysis (**1-3-4**).

#### 1-3-3. Lysis

Each well was washed with 120 µL of cold PBS to remove residual medium. The plate was placed on ice and to each well was added 60 µL of cold RIPA buffer (50 mM Tris-HCl, pH 8.0, 150 mM NaCl, 1% (v/v) Triton X-100, 0.5% (w/v) sodium deoxycholate, 0.1% (w/v) SDS) supplemented with 0.6 µL of 100x protease inhibitor cocktail (Thermo Scientific, 78429) with 250 RPM shaking for 15 minutes. Cell debris was removed from the plate and the cell lysate (60 µL) was combined with 20 µL of 4x SDS-PAGE sample loading buffer (200 mM Tris-HCl, pH 6.5, 400 mM DTT, 8% (w/v) SDS, 6 mM bromophenol blue, 4 M glycerol).

#### 1-3-4. Gel Electrophoresis and Membrane Transfer

Prior to gel electrophoresis, each sample was heated at 95 °C for 10 minutes, separated by 10% (v/v) SDS-PAGE gel electrophoresis (80 V for 20 minutes, 150 V for 80 minutes) with an ice pack placed inside the tank next to the gel cassette. Completed gels were transferred to a 0.45 µm PDVF membrane (Millipore, IPVH00010) at 80 V for 2 hours using cold Transfer Buffer (25 mM Tris-HCl, 192 mM glycine, pH = 8.3, 20% methanol), in an ice bath.

The membrane was blocked for 1 hour with 3 mL of blocking buffer (5% milk in TBS with 0.1% (v/v) Tween 20, TBST) at room temperature with rocking. Blocking buffer was removed and blots were probed with rabbit mAb anti-HA (1:1000 dilution, CST, 3724S) and rabbit pAb anti-GAPDH (1:10000 dilution, ProteinTech, 50-172-6351) primary antibodies in 3 mL of blocking buffer overnight with rocking at 4 °C. After washing three times by incubating the blots with 10 mL of cold TBST with rocking, membranes were incubated for 1 hour with goat anti-rabbit IgG HRP-linked secondary antibody (1:5000 dilution, CST 7074S) in 3 mL of TBST. After washing three times with 3 mL of cold TBST with rocking, the blots were developed in either SuperSignal West Pico Chemiluminescent Substrate (Thermo Scientific, 34580) or homemade imaging solution (Solution A: 39 mg of luminol and 47.5 mg of *p*-iodophenol in 50 mL of 0.1 M Tris-HCl, pH 9.35, and Solution B: 5 µL of 30% hydrogen peroxide in 50 mL of 0.1 M Tris-HCl, pH 9.35). For either chemiluminescent substrate condition, 2 mL of the luminol/enhancer solution was mixed with 2 mL of the peroxide solution and incubating the membrane in the resulting solution at room temperature with rocking for 5 minutes.

#### 1-3-5. Image Analysis

The blots were imaged on a BioRad ChemiDoc system with automated exposure times. Western blots were quantified using ImageJ software: The first individual band/lane was selected using the Rectangular Selection tool, then denoted as "First Lane" through Analyze>Gels>Select First Lane. The newly generated box was positioned over the subsequent lane and denoted as the "Next Lane" through Analyze>Gels>Select Next Lane, and repeated until all bands/lanes were accounted for. Each lane was plotted using Analyze>Gels>Plot Lanes. On the generated graphs, a baseline was added using the Straight Line tool, then the area under the curve was quantified using the Wand tool. The values obtained for each POI were transferred to Excel, then normalized to their respective loading controls (GAPDH), calculated the same way. Then, each POI/GAPDH value was normalized to the average of each respective minus compound controls. Standard error was determined from three biological replicates using one- or two-way ANOVA via Prism 9.

#### 1-3-6. Quantification of Bioconjugation

For the pulse-chase analysis of protein bioconjugation, the initial treatment with the compound of interest (e.g., **1**) will lead to reaction with the TetF-bearing protein of interest to the extent possible. The subsequent treatment with sTCO-CA will lead to reaction with remaining protein not already conjugated by **1**, to the extent determined by treatment with only sTCO-CA. Cell lysis and subsequent incubation with HaloTag will then form a covalent heterodimer with sTCO-CA conjugated protein. Integration of the **non-conjugated** protein bands in the western blot reveals the amount of protein that was conjugated to sTCO-CA. Due to inconsistent western blot transfer efficiencies of proteins with markedly different molecular weights, we do not integrate the band corresponding to HaloTag dimer and have developed the reliable quantification strategy below.

As different proteins and different sites on protein surfaces are conjugated with different efficiencies,[8] their corresponding reactivity needs to be probed first. We achieve this by:

1. Quantify the **non-conjugated** protein from cells treated with DMSO.
2. Quantify **non-conjugated** protein from cells treated with sTCO-CA.
3. Quantify the GAPDH loading control.
4. Normalize the amount of **non-conjugated** protein to its respective GAPDH loading control.

5. Using the values from step 4, normalize the amount of **non-conjugated** protein from each sTCO-CA treatment replicate to the average of **non-conjugated** protein from DMSO treated cells.
6. The average of the three replicates equals the amount of protein that cannot be conjugated by sTCO-CA. Subtract this number from 1 to determine the **bioconjugation efficiency** of sTCO-CA.

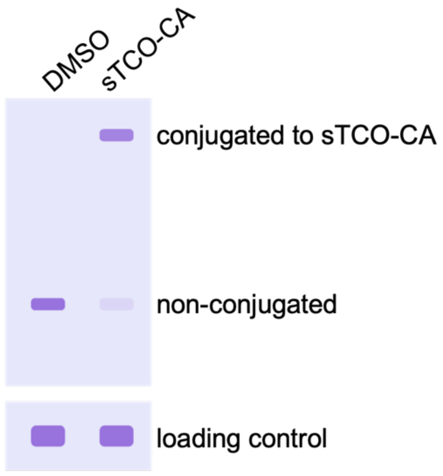

The **bioconjugation efficiency** of sTCO-CA will then be used to calculate the protein bioconjugation yield by our ligand **1** via pulse-chase (P-C) analysis:

1. Quantify **non-conjugated** protein from cells treated with **1** only.
2. Quantify **conjugated to 1 or non-conjugated** protein from cells treated with **1** then treated with sTCO-CA.
3. Normalize the values from steps 1 and 2 to the density of the respective GAPDH loading controls.
4. Using the GAPDH-normalized values from step 3 normalize the bottom pulse-chase band for each replicate to the average of the bands from cells treated with **1** only. This will indicate **sTCO-CA unreacted protein**.
5. Determine the amount of non-conjugated protein remaining bymultiplying the **sTCO-CA unreacted protein** by the **bioconjugation efficiency** calculated above for each mutant.

#### 1-4. siRNA knockdown of CRBN

HEK293T cells were seeded into a poly-D-lysine (PDK) treated 48-well plate (50,000 cells/well, Greiner, 677180) in 250 μL of complete medium. When cells reached 60-70% confluency, each well was washed once with 200 μL of OptiMEM, then replaced with another fresh aliquot of 172 μL of fresh OptiMEM. The plate was returned to the incubator for the preparation of the transfection solution.

To transfect cells in one well with 100 nM of CRBN siRNA (human, SCBT sc-78528), 2 µL of siRNA (10 µM in RNAse-free water) was diluted into 12.5 µL of OptiMEM, and labeled as Tube A. In a separate microcentrifuge tube, Tube B, 0.75 µL of Lipofectamine3000 (Fisher L3000008), 0.5 µL of P3000 reagent (Fisher L3000008), and 12.5 µL of OptiMEM were combined. Tube B was pipetted to mix, then combined with the mixture in Tube A. The resultant transfection mixture was incubated at room temperature for 15 min. Then, 28 µL of the transfection mixture was added dropwise to each well. For cells not transfected with siRNA, the same protocol was followed, excluding the addition of siRNA to Tube A. The cells were incubated for 12 hours at 37 °C and 5% $CO_2$, then stimulated with antibiotic-free DMEM supplemented with 250 µM of **TetF**, containing a final DMSO concentration of 0.25%.

After stimulation, the cells were then transfected cells with the necessary genetic code expansion machinery (pAcBac-TetFRS and pmCherry-EGFP-Y151TAG-HA) using LPEI, as described above. Briefly, for each well, 400 ng of DNA (1:2 TAG-mutant/pAcBac-TetFRS) and 2 µL of LPEI were diluted into 40 µL of OptiMEM transfection media. The transfection master mix was gently mixed by inversion and incubated for 10 minutes at room temperature. Finally, 40 µL of the mixture was added dropwise to each well and the cells were then incubated at 37 °C and 5% $CO_2$ for 24 hours.

Following overnight transfection, the cells were washed by gently applying 200 µL of fresh complete medium to each well and immediately removing the media. A second wash was completed by adding another 200 µL of fresh complete medium to each well and incubating the cells at 37 °C for 30 minutes before gently removing the supernatant. After UAA washout, cells were then treated with or without 10 µM of **1a** overnight. Cells were then lysed, as described in section 1-3-3, and lysate was analyzed via western blot (section 1-3-4). Anti-CRBN western blots were performed using rabbit anti-CRBN (Sigma HPA045910) at a 1:1000 dilution in 5% milk in TBST, incubated at 4 °C with rocking overnight.

#### 1-5. Docking

The small molecule ligand BCI was docked into DUSP6 (PDB 1MKP) according to literature protocols[10] using Chimera and Autodock Vina. The protein structure of DUSP6 was retrieved from the RCSB protein database using its PDB code. The structure was prepared for docking using Tools>Structure Editing>Dock Prep. In the new window, the following options were selected: Delete solvent, If alternate locations, keep only highest occupancy, Incomplete side chains: Replace using Dunbrack rotamer library, Add hydrogens, Add charges, and Write Mol2 file. In the next “Add Hydrogens” window, no parameters were changed. In the next, “Assign Charges for Dock Prep” window, Gasteiger charges were selected. No changes were made in the following window, “Specify Net Charges.” The prepared structure was saved in a separate file. The BCI ligand was then prepared in Chimera under Tools>Structure Editing>Build Structure. The SMILES code (O=C(/C(C1NC2CCCCC2)=C/C3=CC=CC=C3)C4=C1C=CC=C4) was added and applied to the structure. To then dock BCI into the region outlined in the literature, Tools>Surface/Binding Analysis>Autodock Vina. Under “Receptor search volume options” the box parameters were set to Center: 6, 54, 5 and Size: 12, 13, 10. No other parameters in Autodock Vina were changed.

### 2. Synthetic Protocols

#### 2-1. General Chemical Methods

All reagents and solvents were purchased from commercial suppliers and used without further purification. All reactions were stirred magnetically and flash chromatography on silica gel (standard grade, 60A, 40-63 µm, Sorbtech, 40930H-25) was performed by hand. NMR spectra were recorded on Bruker Ultrashield 400 MHz or 500 MHz spectrometers. Analytical LC-MS data were collected on a Shimadzu LCMS-2020 and a Thermo Scientific Q-Exactive Orbitrap. High resolution mass spectrometry was performed by the University of Pittsburgh facilities.

#### 2-2. Synthesis of sTCO-PEG$_n$-CRBN

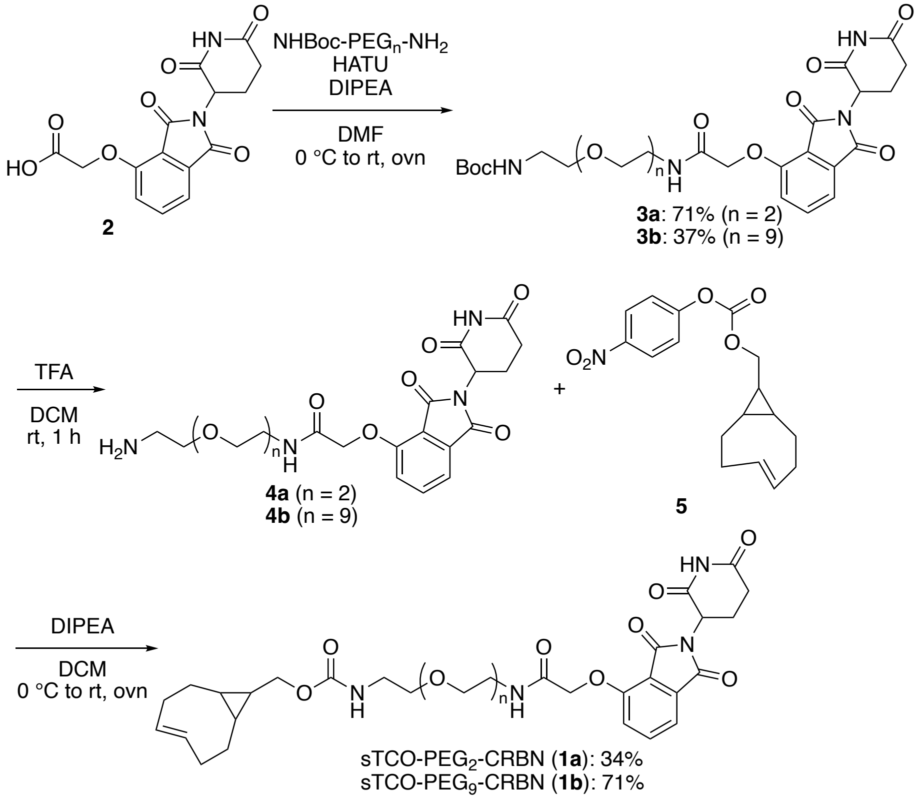

#### General Procedure

Compound **2** was synthesized and characterized according to the literature.[11] In a flame-dried vial, the carboxylic acid **2** (1 eq) was dissolved in DMF (final carboxylic acid concentration of 0.1 M), then HATU (1.1 eq) and DIPEA (4 eq) were added, and the mixture was stirred on ice for 5 minutes.[11] The NHBoc-PEG$_n$-Amine (1.1 eq) was added and the mixture was allowed to warm to room temperature overnight. Following consumption of the carboxylic acid as observed on TLC

(5% MeOH/DCM), the reaction was diluted with brine (10 times the volume of DMF used in the reaction) and extracted four times with EtOAc (1-10 mL). The combined organic layers were dried over sodium sulfate and concentrated *in vacuo*. The resultant crude oil was subjected to normal phase flash chromatography using 5% MeOH/DCM to elute the product **3**.

Following characterization of the NHBoc-PEG$_n$-CRBN product, the material was dissolved in 500 µL of 25% TFA in DCM and stirred at room temperature open to the air.[11] When complete consumption of the Boc-protected starting material was observed by TLC (10% MeOH/DCM, usually 30-60 minutes), the reaction was concentrated *in vacuo*. Excess TFA was removed by resuspending the crude material in 2 mL of methanol and concentrating to dryness, then repeating this process 3 times. Crude **4** was characterized by LC-MS and then directly used in the next step without further purification.

Compound **5** was synthesized and characterized according to the literature,[5] then added as a neat solid (1.2 eq) to a solution of the amine **4** in DCM (final amine concentration of 0.05 M) and DIPEA (4 eq) in a flame-dried vial. The reaction vial was wrapped in aluminum foil and the mixture was stirred at room temperature overnight.[12,13] The reaction was then concentrated *in vacuo* and the crude material was subjected to normal phase flash chromatography (5 to 10% MeOH/DCM). The resultant material collected contained 4-nitrophenol impurities, which were not separable by flash chromatography. To remove this impurity, the mixture was dissolved in DCM (1 mL/50 mg) and extracted with small amounts of saturated sodium carbonate (2-5 mL).[12] The organic layer was concentrated *in vacuo* and the resultant, pure product was characterized by $^1$H NMR, $^{13}$C NMR, LC-MS, and HRMS.

**NHBoc-PEG$_2$-CRBN (3a).** The following reactants and reagents yielded 237 mg of **3a** (71% yield) as a white solid: **2** (196 mg, 0.59 mmol), BocNH-PEG$_2$-NH$_2$ (160 mg, 0.64 mmol), HATU (247 mg, 0.65 mmol), DIPEA (200 µL, 1.8 mmol), and DMF (5.9 mL). $^1$H NMR (400 MHz, CDCl$_3$) δ 8.93 (bs, 1H), 7.73 (t, 1H, *J* = 7.6 Hz), 7.62 (bs, 1H), 7.53 (d, 1H, *J* = 7.2 Hz), 7.18 (d, 1H, *J* = 8.4 Hz), 5.15 (bs, 1H), 4.96 (m, 1H), 4.66 (s, 2H), 3.58 (m, 10 H), 3.28 (bs, 2H), 2.81 (m, 3H), 2.15 (m, 1H), 1.44 (s, 9H). $^{13}$C NMR (CDCl$_3$, 100 MHz) 174.7, 171.4, 168.5, 167.1, 166.8, 166.0, 156.3, 154.6, 137.1, 133.8, 119.5, 118.1, 117.4, 70.4, 70.1, 69.6, 68.0, 53.6, 49.4, 39.2, 38.9, 36.7, 31.6, 29.8, 28.5, 22.8, 20.8.

**NHBoc-PEG$_9$-CRBN (3b)**. The following reactants and reagents yielded 58 mg of **3b** (37% yield) as a clear oil: **2** (70 mg, 0.21 mmol), NHBoc-PEG$_9$-NH$_2$ (100 mg, 0.18 mmol), HATU (94 mg, 0.25 mmol), DIPEA (150 µL, 0.8 mmol), and DMF (2 mL). $^1$H NMR (500 MHz, CDCl$_3$) δ 8.89 (bs, 1H), 7.75 (t, 1H, *J* = 7.6 Hz), 7.68 (m, 1H), 7.54 (d, 1H, *J* = 4.8 Hz), 7.22 (d, 1H, *J* = 8.4 Hz), 6.14 (bs, 1 H), 4.95 (m, 1H), 4.66 (s, 2H), 3.60 (m, 32H), 3.52 (m, 4H), 3.38 (m, 2H), 3.31 (m, 4H), 3.18 (m, 2H), 3.0 (bs), 2.16 (m, 1H), 1.5 (s, 9H). $^{13}$C NMR (125 MHz, CDCl$_3$) 171.4, 168.5, 167.4, 166.8, 166.0, 156.4, 154.6, 137.2, 133.7, 119.8, 118.2, 117.4, 70.7, 70.3, 70.2, 70.1, 70.0, 69.9, 69.8, 69.8, 69.3, 68.1, 54.9, 53.6, 49.4, 40.4, 39.1, 31.5, 29.8, 28.5, 22.7, 21.5, 18.7, 17.3, 12.5. HRMS $(M+H)^+$ calcd for $C_{40}H_{62}N_4O_{17}$ $(M+H)^+$ 870.9470, found 871.4201.

**sTCO-PEG$_2$-CRBN** (**1a**). The following reactants and reagents yielded 35 mg of **1a** (34% yield) as a white solid: **4a** (100 mg, 0.22 mmol), **5** (50 mg, 0.16 mmol), DIPEA (110 µL, 0.64 mmol), and

DCM (20 mL). $^1$H NMR (500 MHz, $CDCl_3$) δ 8.68 (bs, 1H), 7.73 (t,1H, *J* = 7.5 Hz), 7.61 (m, 1H), 7.55 (d, 1H, *J* = 7 Hz), 7.19 (d, 1H, *J* = 8.5 Hz), 6.85 (m, 1H), 5.35 (m, 1H), 5.11 (1 H, m), 4.96 (dd, 1H, J = 5.5, 5), 4.66 (s, 2H), 3.93 (d, 2H, *J* = 6 Hz), 3.60 (m, 11H), 3.34 (s, 2H), 2.83 (m, 3H), 2.14 (m, 8 H), 1.42 (m, 1H), 1.26 (m, 5H), 0.85 (m, 2H), 0.52 (m, 2H), 0.40 (m, 2H). $^{13}$C NMR (125 MHz, $CDCl_3$) 171.2, 168.3, 167.1, 166.8, 166.0, 157.2, 154.6, 138.5, 137.1, 133.8, 131.4, 126.3, 119.5, 118.2, 117.5, 115.8, 70.4, 70.3, 70.2, 69.7, 68.1, 53.6, 49.4, 40.9, 39.2, 38.8, 33.9, 32.7, 31.5, 29.8, 29.7, 29.5, 29.4, 27.8, 24.8, 22.8, 22.1, 21.1, 14.3. HRMS $(M+H)^+$ calcd for $C_{32}H_{40}N_4O_{10}$ 640.2744, found 639.2639.

**sTCO-PEG$_9$-CRBN** (**1b**). The following reactants and reagents yielded 15 mg of **1b** (71% yield) as a clear oil: **4b** (22 mg, 0.03 mmol), **5** (9.5 mg, 0.03 mmol), DIPEA (20 µL, 0.12 mmol), and DCM (600 µL). $^1$H NMR (400 MHz, $CDCl_3$) δ 8.95 (bs, 1H), 7.75 (t, 1H, *J* = 7.5 Hz), 7.65 (m, 1H), 7.55 (d, 1H, *J* = 7 Hz), 5.86 (m, 1H), 5.11 (m, 1H), 4.65 (s, 1H), 3.93 (s, 2H), 3.65 (m, 32H), 3.57 (m, 4H), 3.35 (s, 2H), 2.78 (m, 4H), 2.25 (m, 6H), 1.91 (m, 6H), 0.85 (m, 2H), 0.54 (m, 2H), 0.42 (m, 2H). $^{13}$C NMR (100 MHz, $CDCl_3$) 171.2, 168.2, 166.9, 166.8, 165.9, 157.0, 154.6, 138.5, 137.1, 133.8, 131.4, 119.5, 118.2, 117.4, 70.75, 70.70, 70.67, 70.64, 70.59, 70.58, 70.55, 70.48, 70.44, 70.3, 69.6, 69.5, 68.0, 49.5, 40.9, 39.2, 38.8, 33.9, 32.7, 31.6, 27.8, 24.9, 22.8, 22.1, 21.1. HRMS $(M+H)^+$ calcd for $C_{46}H_{68}N_4O_{17}$ $(M+H)^+$ 948.4579, found 947.4507.

### 3. Supplemental Figures and Tables

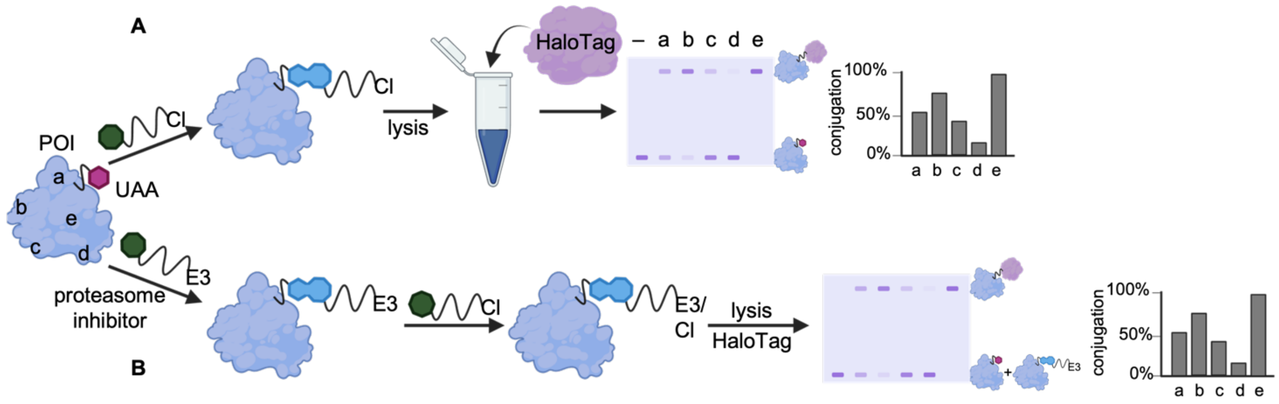

**Figure S1. Experimental workflow for the quantification of bioconjugation to UAA-bearing proteins expressed in cells.**

Following expression of the POI bearing a UAA at any surface position (e.g., A-E), the A) sTCO-chloroalkane ligand or B) sTCO-E3 ligase ligand, is added to the media and undergoes a selective bioconjugation with the UAA. Excess ligand is washed away and cells are A) lysed or B) incubated with sTCO-chloroalkane then lysed. The lysate is incubated with HaloTag protein and the bioconjugation is quantified via western blot.

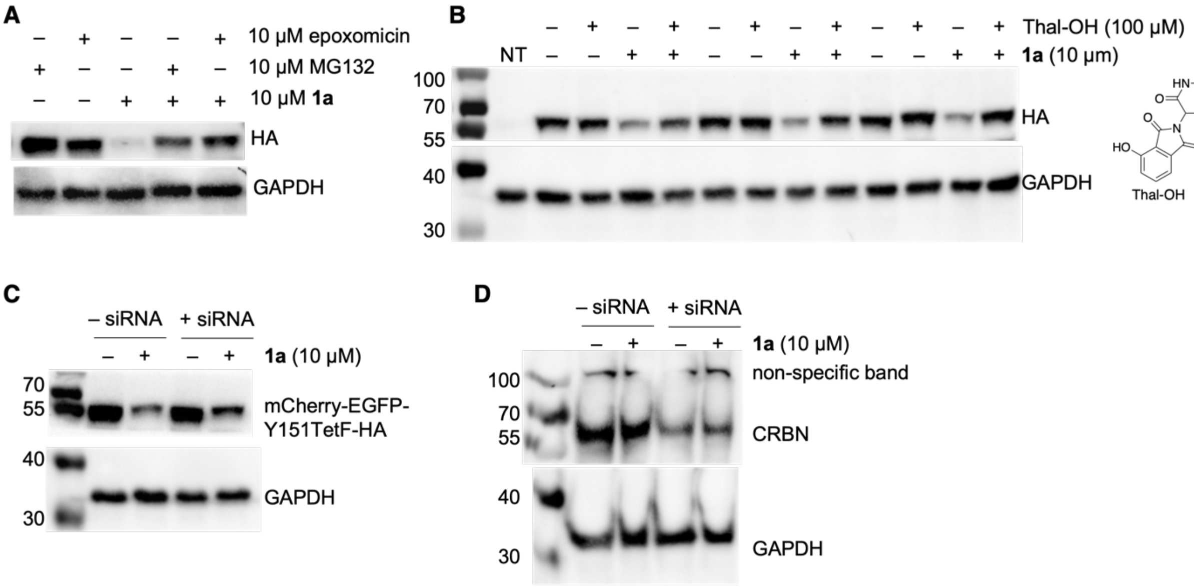

**Figure S2. Degradation mechanism of 1a.**

A) The degradation of mCherry-EGFP-Y151TetF was reduced when cells were pre-treated with proteasome inhibitor MG132 or epoxomicin, validating that **1a** induced degradation is mediated by the proteasome. B) Pretreatment of cells expressing mCherry-EGFP-Y151**TetF** with 100 μM of thalidomide-OH (Thal-OH) followed by treatment with 1a reduced degradation, validating cereblon-mediated degradation. C) Co-transfection of HEK293T cells with CRBN-siRNA and genetic code expansion machinery decreased the degradation of mCherry-EGFPY151**TetF**-HA in the presence of **1a**. D) Co-transfection of HEK293T cells with CRBN-siRNA and genetic code expansion machinery decreased the amount of cereblon expressed in cells. Each method validates the cereblon-mediated degradation mechanism of **1a**.

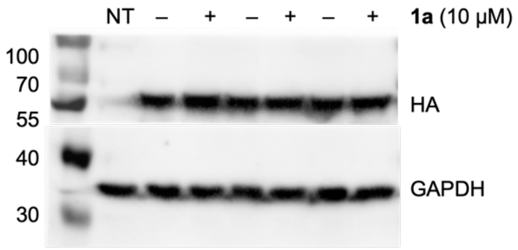

**Figure S3. Specificity of 1a for TetF.**

No background reactivity was observed upon treatment of wild-type mCherry-EGFP-HA with **1a**.

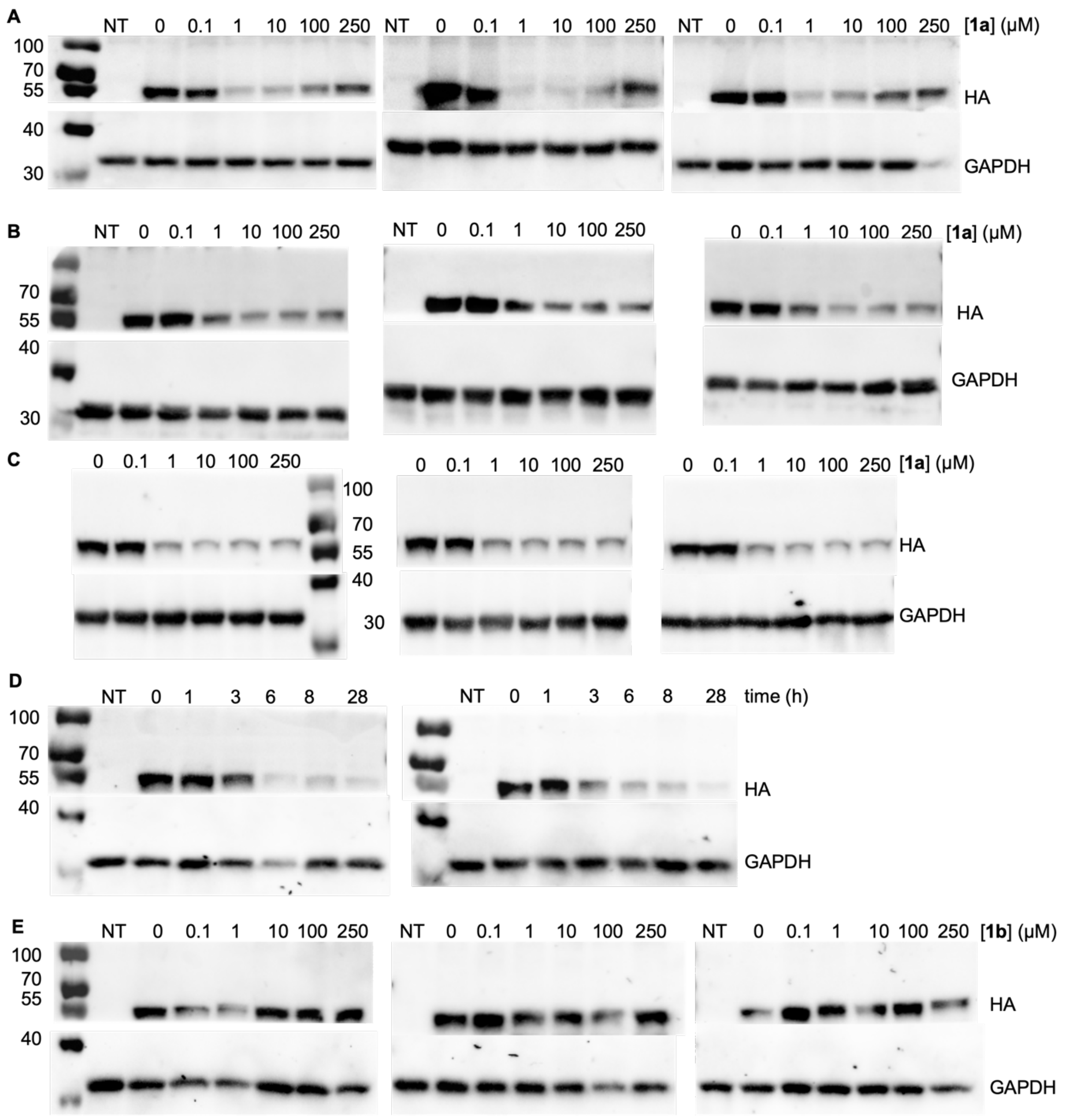

**Figure S4. Full HA and GAPDH blots for each quantification shown in Figure 2.**

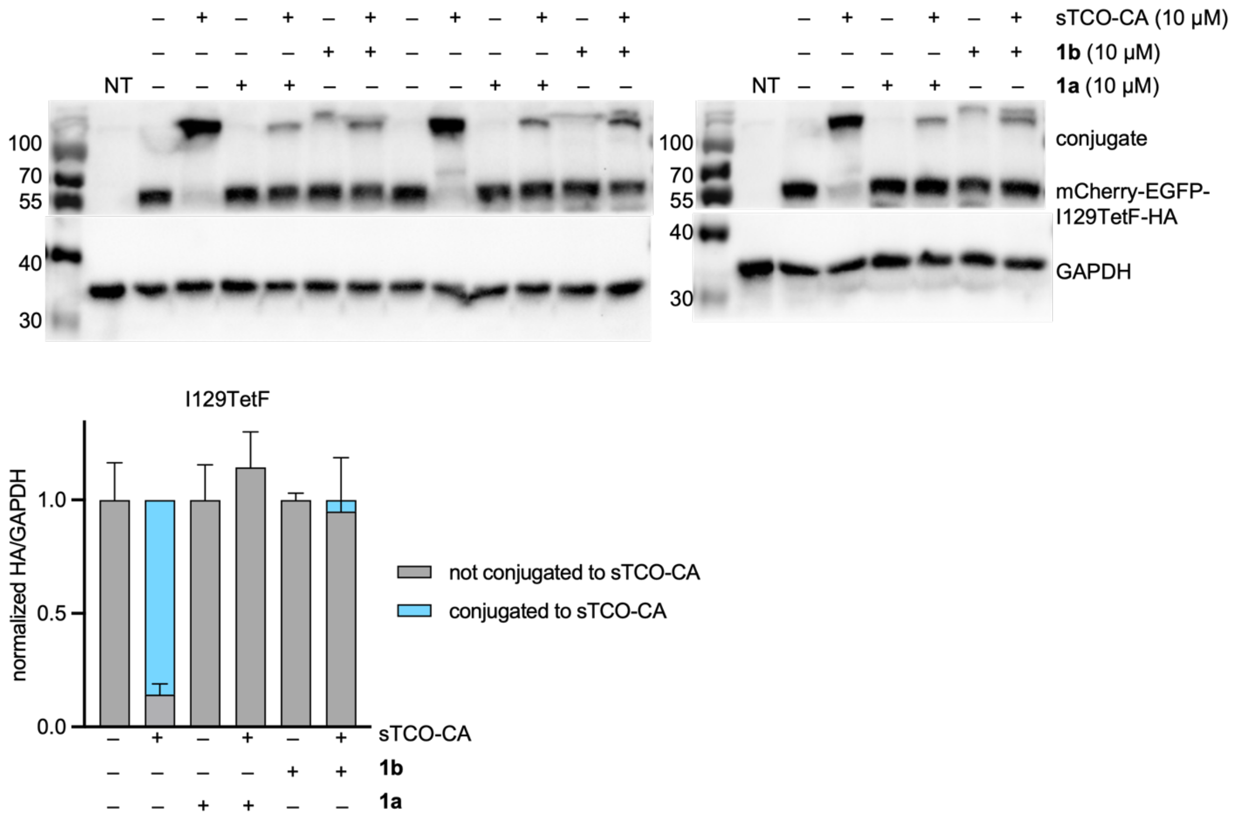

**Figure S5. Pulse-Chase Quantification of Bioconjugation for mCherry-EGFP-I129TetF-HA.**

The bioconjugation efficiency of sTCO-CA, **1a**, and **1b** as determined via triplicate pulse-chase experiment in the presence of the proteasome inhibitor MG132 (10 µM).

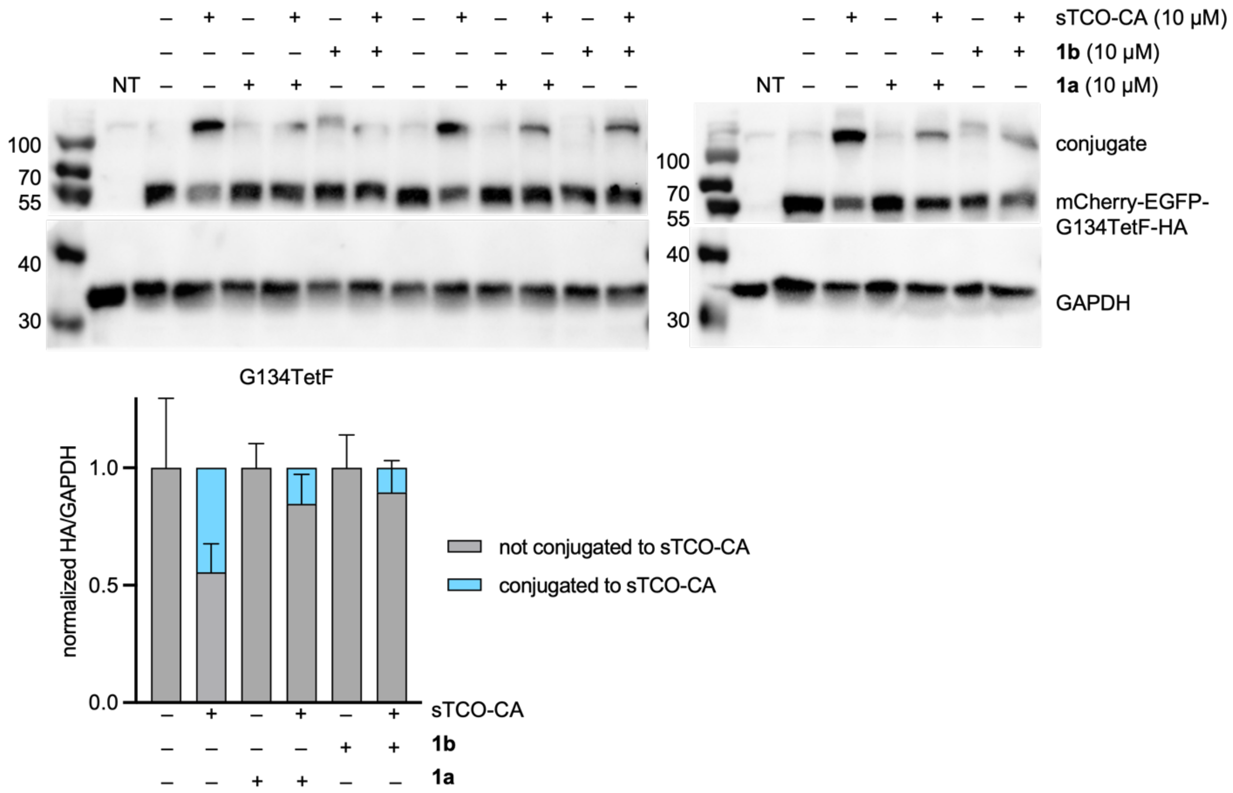

**Figure S6. Pulse-Chase Quantification of Bioconjugation for mCherry-EGFP-G134TetF-HA.**

The bioconjugation efficiency of sTCO-CA, **1a**, and **1b** as determined via triplicate pulse-chase experiment in the presence of the proteasome inhibitor MG132 (10 μM).

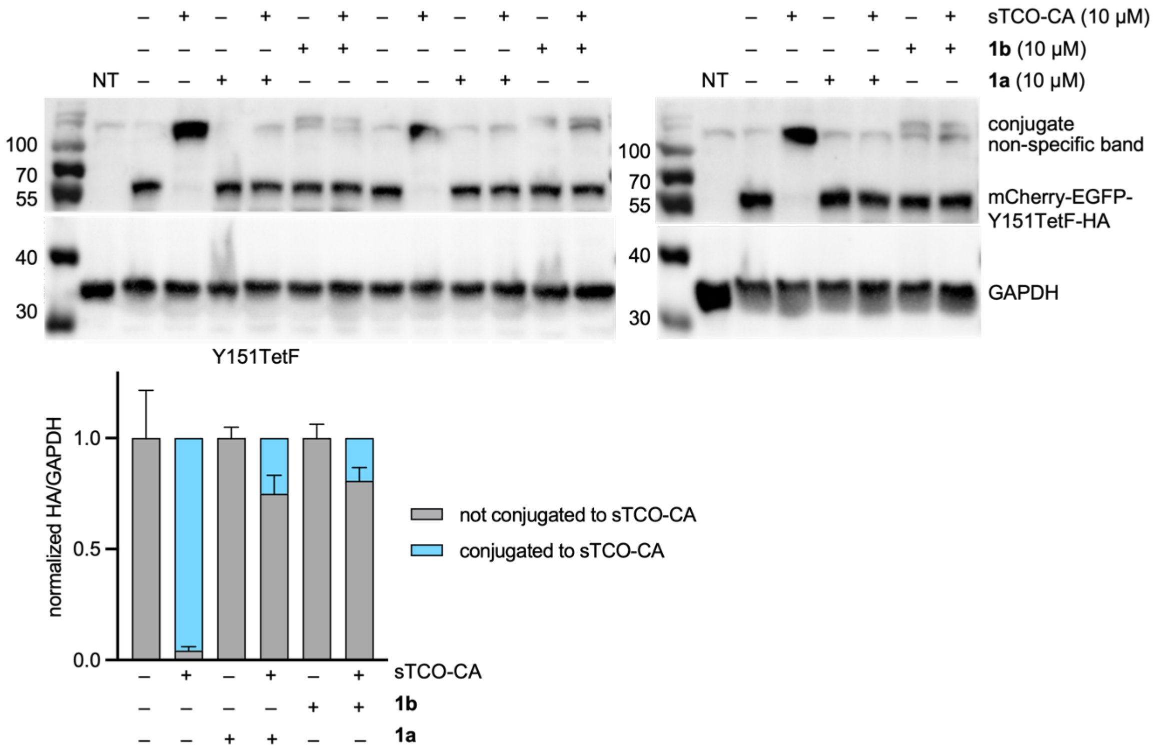

**Figure S7. Pulse-Chase Quantification of Bioconjugation for mCherry-EGFP-Y151TetF-HA.**

The bioconjugation efficiency of sTCO-CA, **1a**, and **1b** as determined via triplicate pulse-chase experiment in the presence of the proteasome inhibitor MG132 (10 μM).

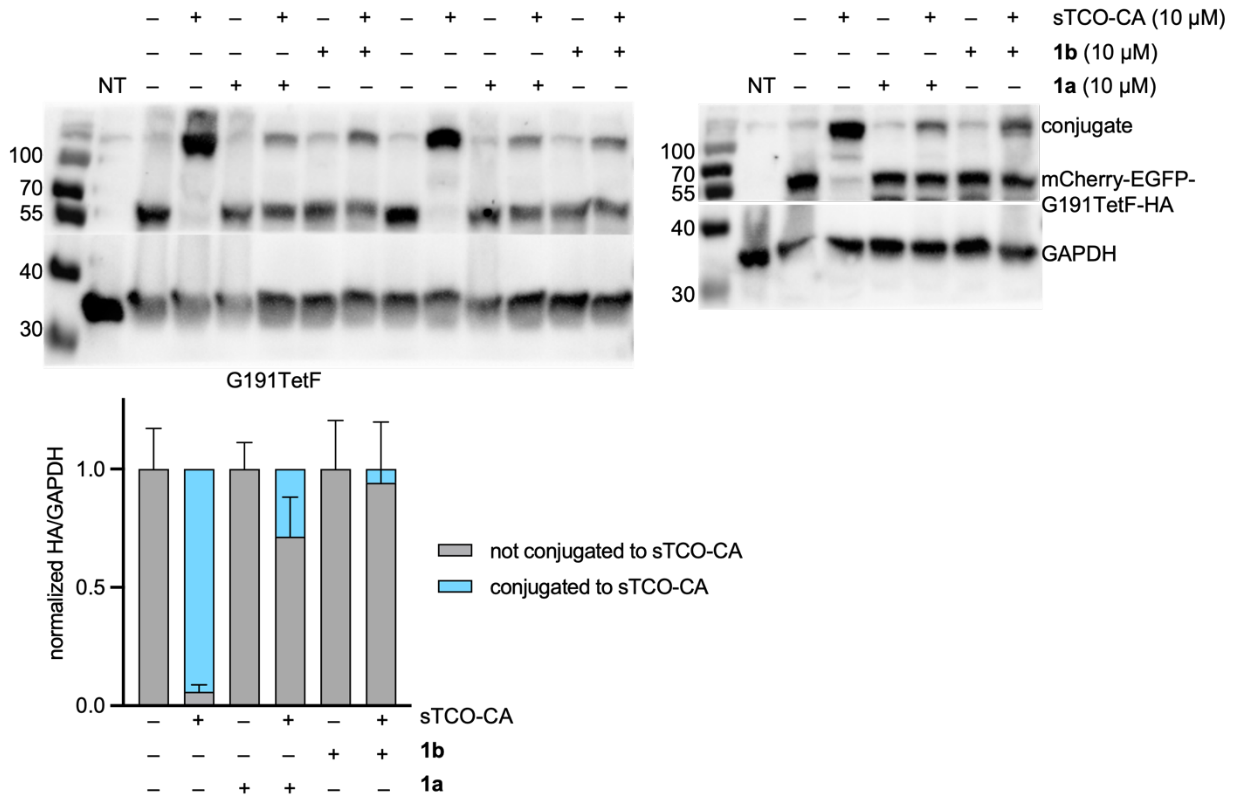

**Figure S8. Pulse-Chase Quantification of Bioconjugation for mCherry-EGFP-G191TetF-HA.**

The bioconjugation efficiency of sTCO-CA, **1a**, and **1b** as determined via triplicate pulse-chase experiment in the presence of the proteasome inhibitor MG132 (10 µM).

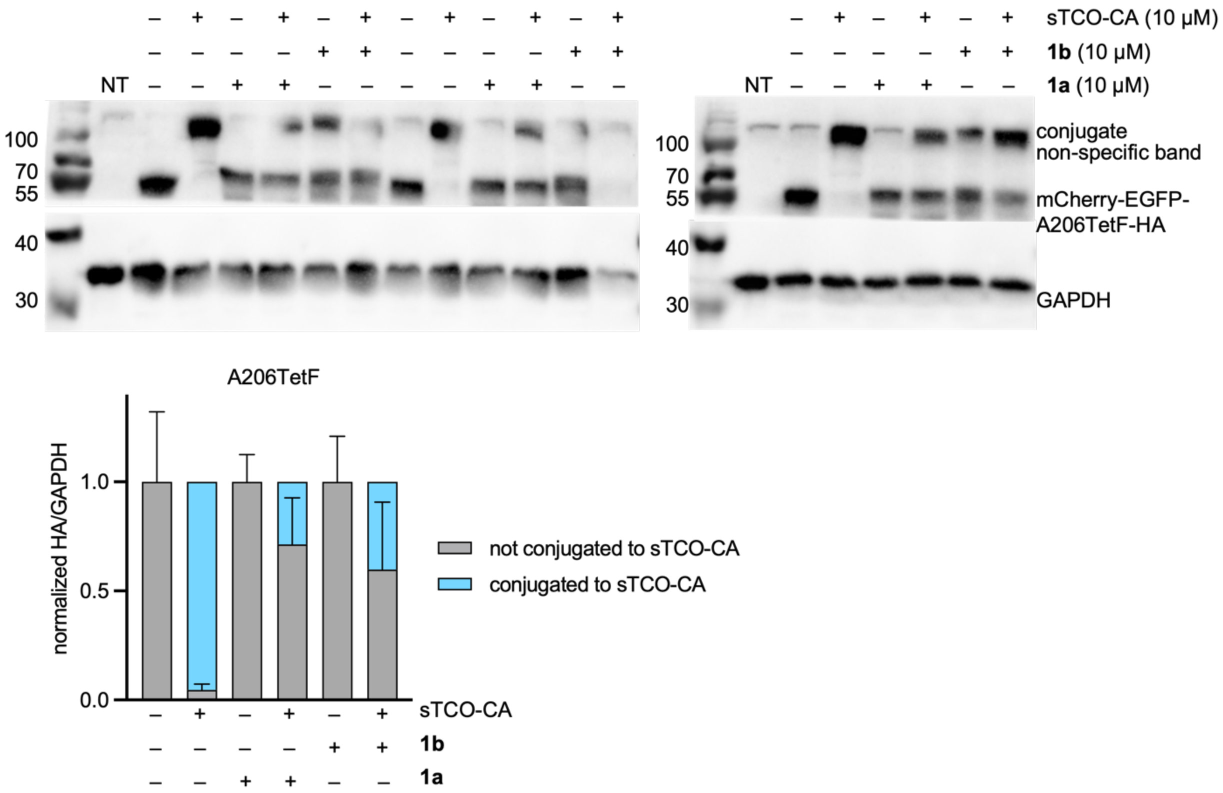

**Figure S9. Pulse-Chase Quantification of Bioconjugation for mCherry-EGFP-A206TetF-HA.**

The bioconjugation efficiency of sTCO-CA, **1a**, and **1b** as determined via triplicate pulse-chase experiment in the presence of the proteasome inhibitor MG132 (10 µM).

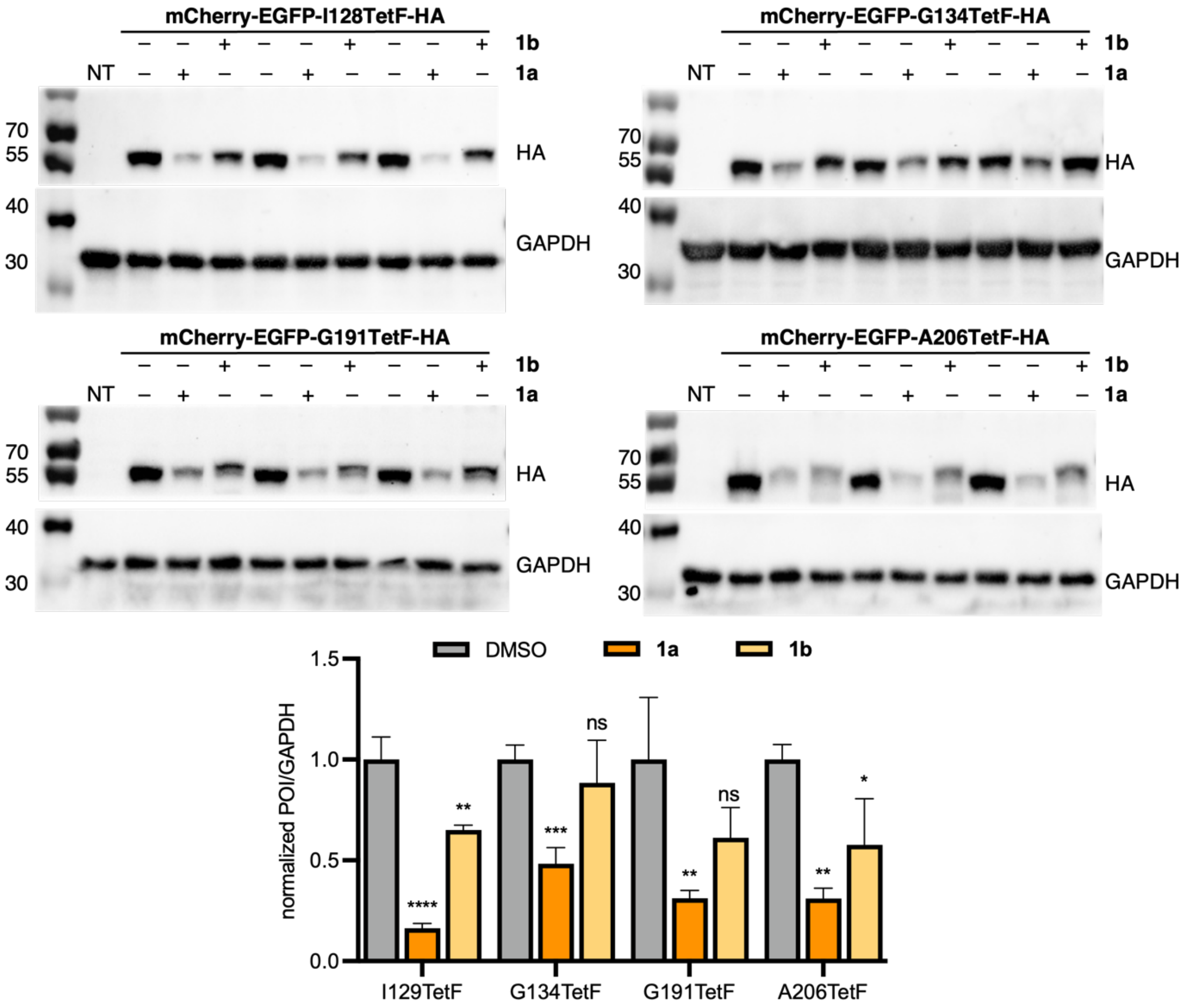

**Figure S10. Full HA and GAPDH blots for each quantification shown in Figure 3B-C.**

Statistical analysis of three biological replicates. Error bars were calculated from three biological replicates, with statistical significance determined through one-way ANOVA, where ** = $p < 0.01$, *** = $p < 0.001$, and **** = $p < 0.0001$.

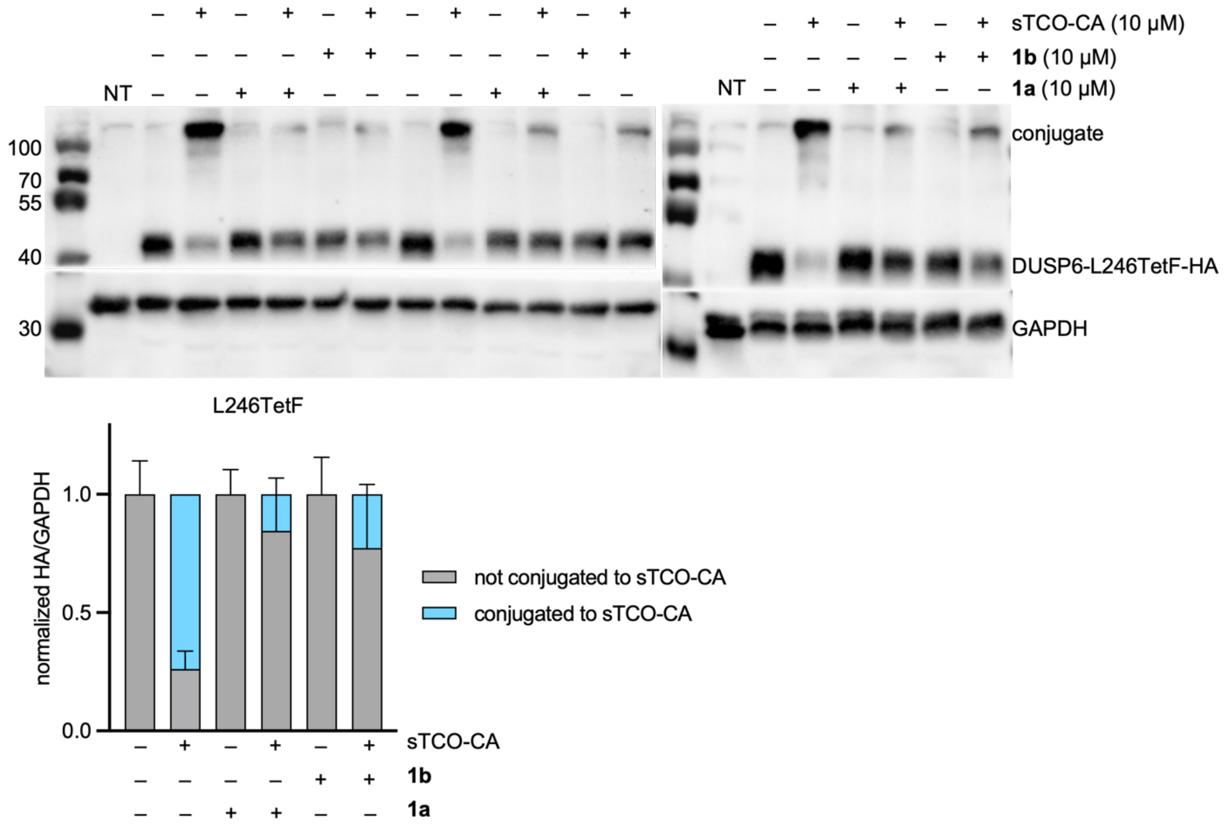

**Figure S11. Pulse-Chase Quantification of Bioconjugation for DUSP6-L246TetF-HA.**

The bioconjugation efficiency of sTCO-CA, **1a**, and **1b** as determined via triplicate pulse-chase experiment in the presence of the proteasome inhibitor MG132 (10 μM).

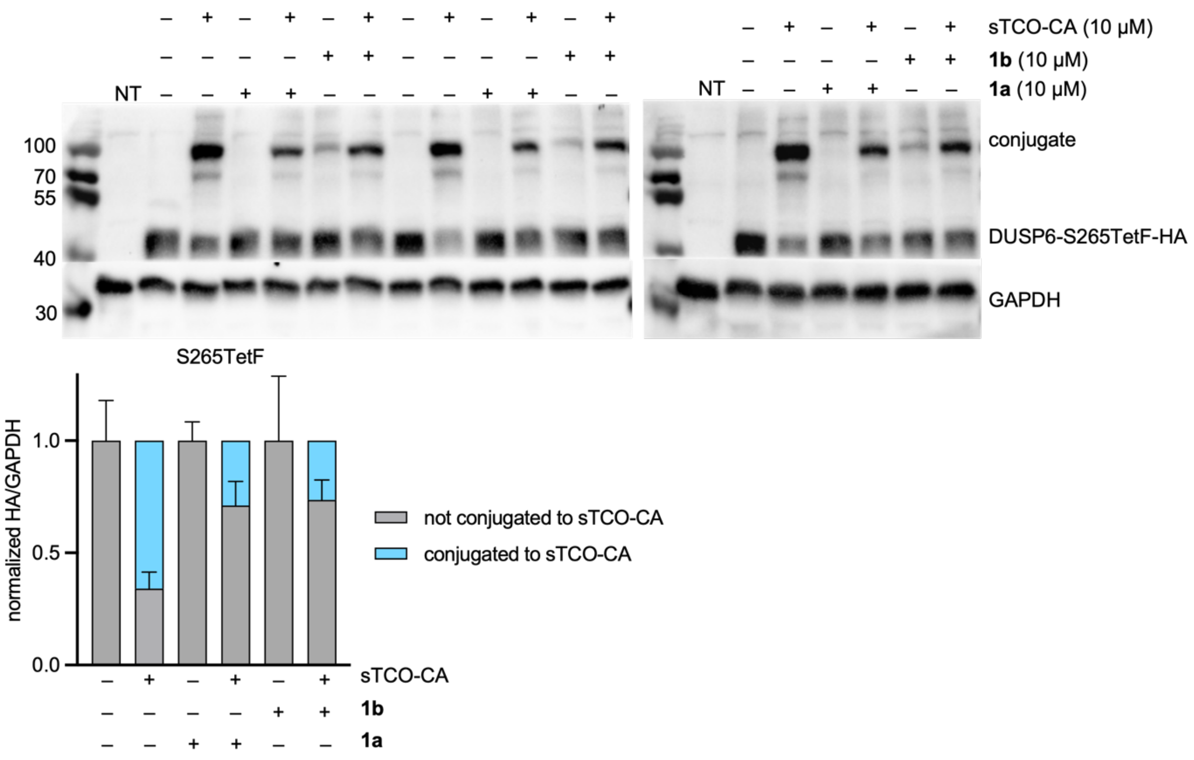

**Figure S12. Pulse-Chase Quantification of Bioconjugation for DUSP6-S265TetF-HA.**

The bioconjugation efficiency of sTCO-CA, **1a**, and **1b** as determined via triplicate pulse-chase experiment in the presence of the proteasome inhibitor MG132 (10 µM).

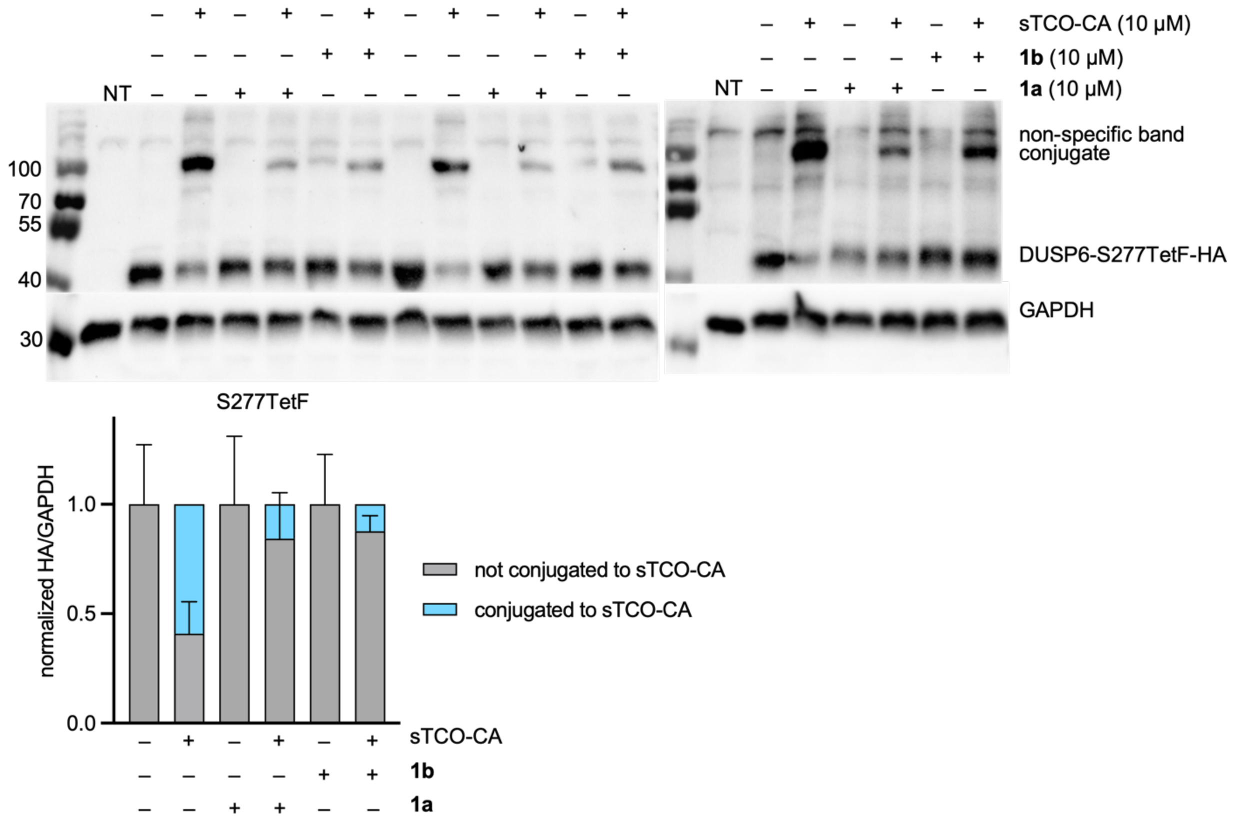

**Figure S13. Pulse-Chase Quantification of Bioconjugation for DUSP6-S277TetF-HA.**

The bioconjugation efficiency of sTCO-CA, **1a**, and **1b** as determined via triplicate pulse-chase experiment in the presence of the proteasome inhibitor MG132 (10 μM).

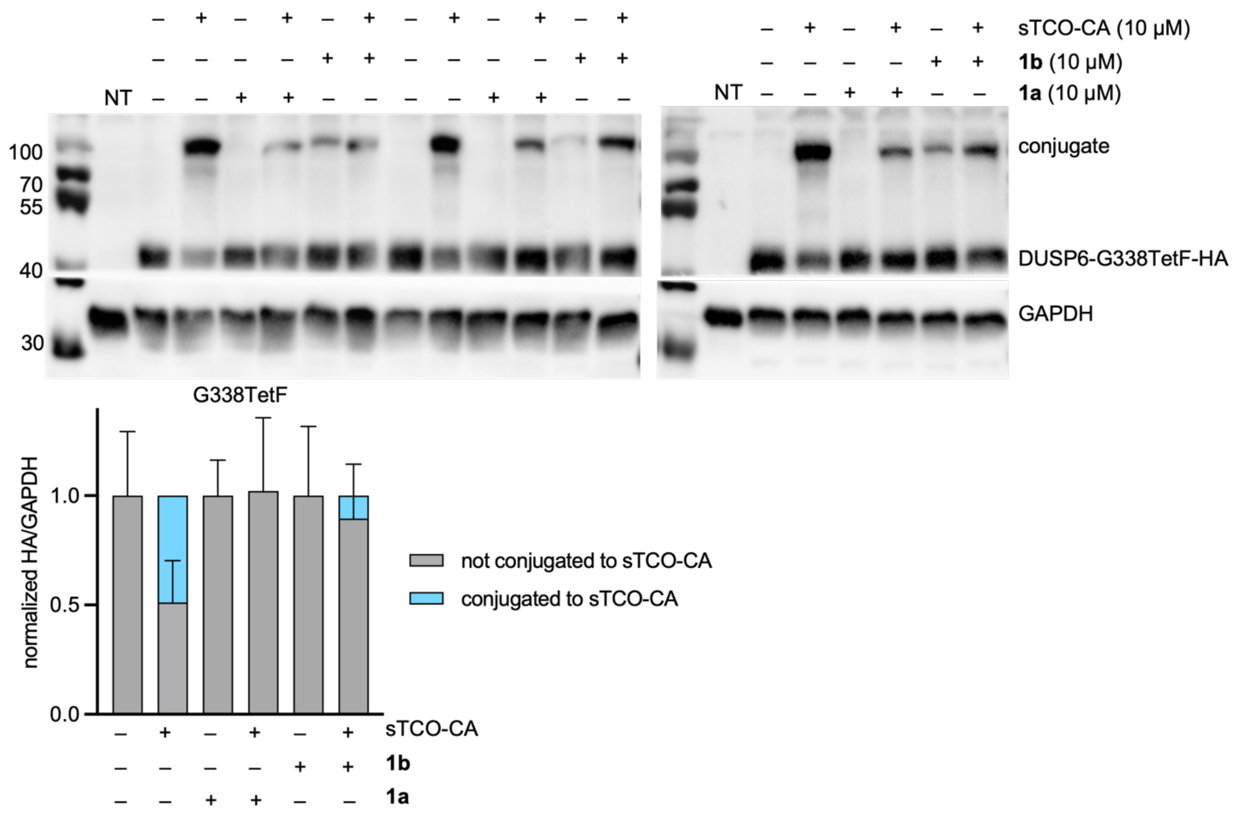

**Figure S14. Pulse-Chase Quantification of Bioconjugation for DUSP6-G338TetF-HA.**

The bioconjugation efficiency of sTCO-CA, **1a**, and **1b** as determined via triplicate pulse-chase experiment in the presence of the proteasome inhibitor MG132 (10 μM).

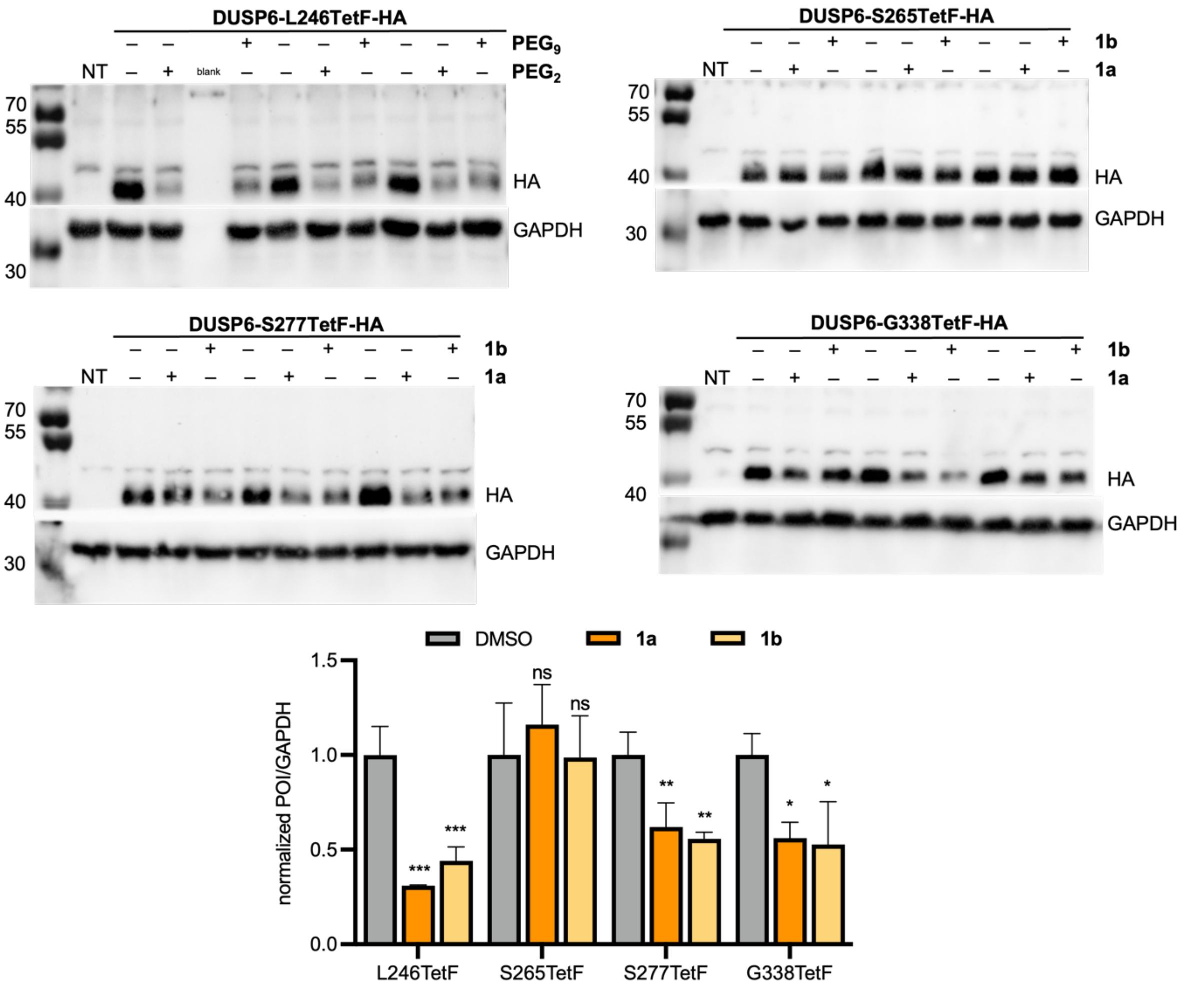

**Figure S15. Full HA and GAPDH blots for each quantification shown in Figure 4B.**

Error bars were calculated from three biological replicates, with statistical significance determined through one-way ANOVA or paired t-test, where *** = $p < 0.001$, and **** = $p < 0.0001$.

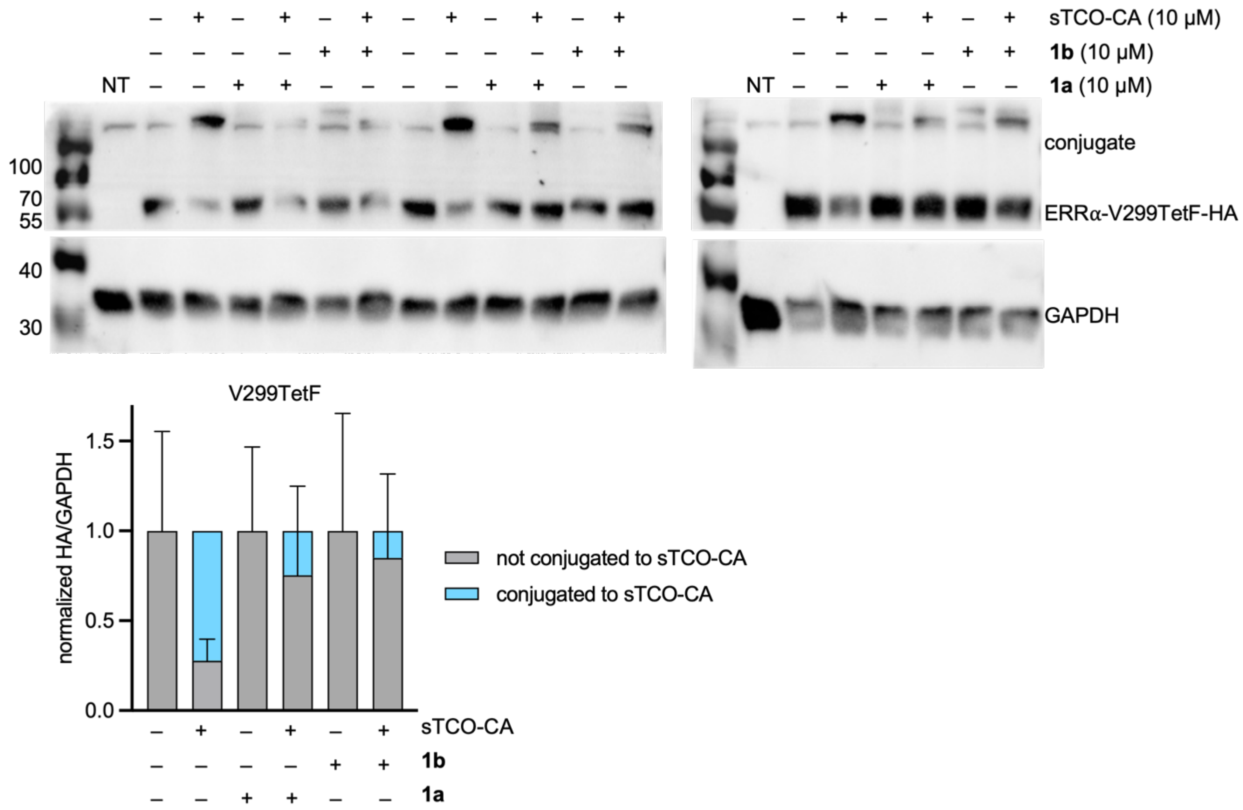

**Figure S16. Pulse-Chase Quantification of Bioconjugation for ERRα-V299TetF-HA.**

The bioconjugation efficiency of sTCO-CA, **1a**, and **1b** as determined via triplicate pulse-chase experiment in the presence of the proteasome inhibitor MG132 (10 μM).

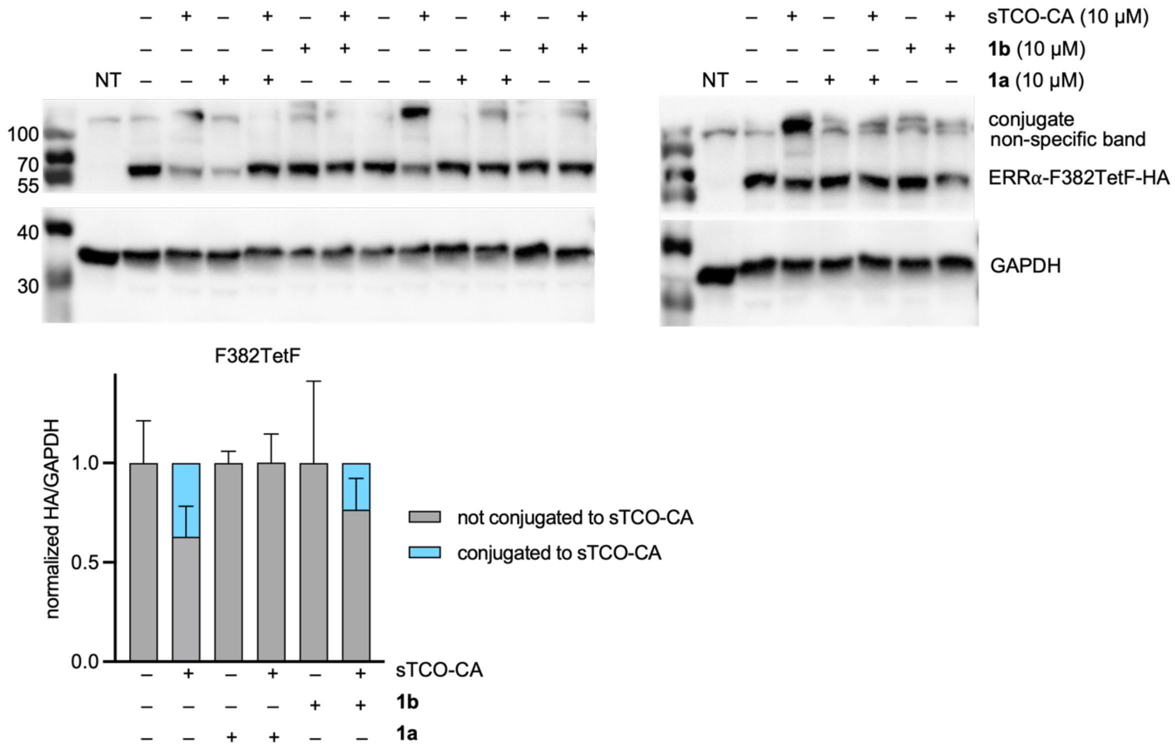

**Figure S17. Pulse-Chase Quantification of Bioconjugation for ERRα-F382TetF-HA.**

The bioconjugation efficiency of sTCO-CA, **1a**, and **1b** as determined via triplicate pulse-chase experiment in the presence of the proteasome inhibitor MG132 (10 μM).

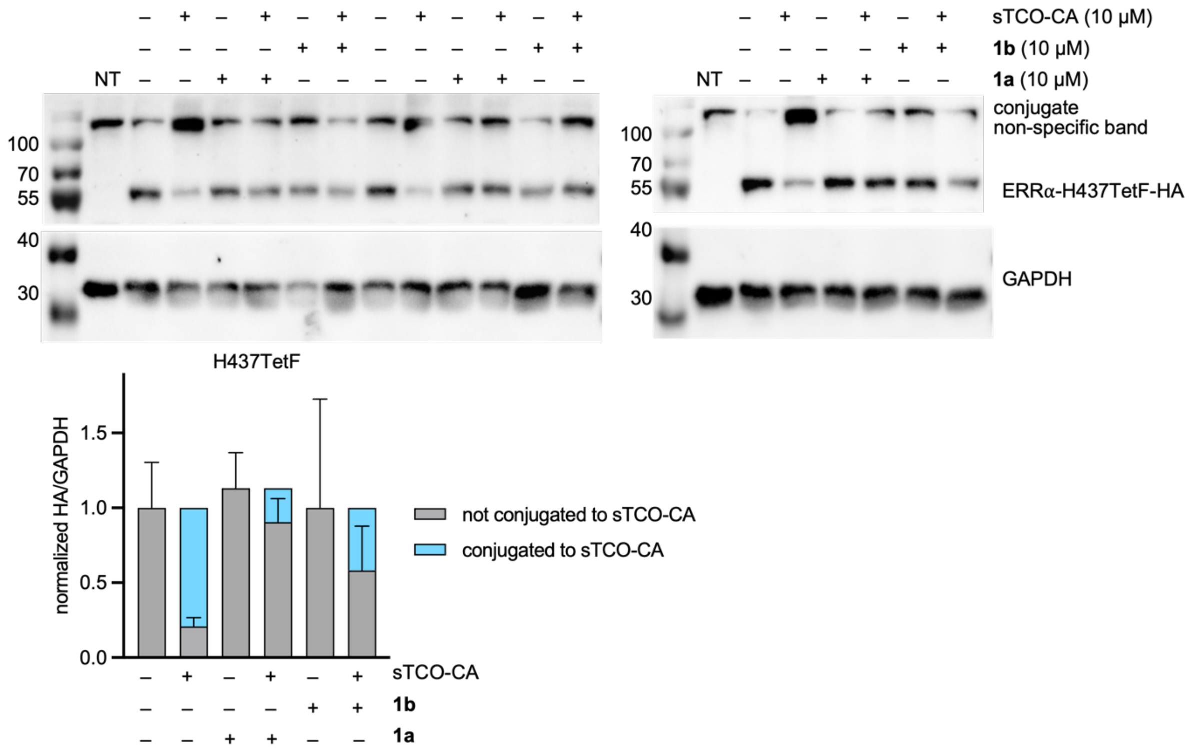

**Figure S18. Pulse-Chase Quantification of Bioconjugation for ERRα-H437TetF-HA.**

The bioconjugation efficiency of sTCO-CA, **1a**, and **1b** as determined via triplicate pulse-chase experiment in the presence of the proteasome inhibitor MG132 (10 μM).

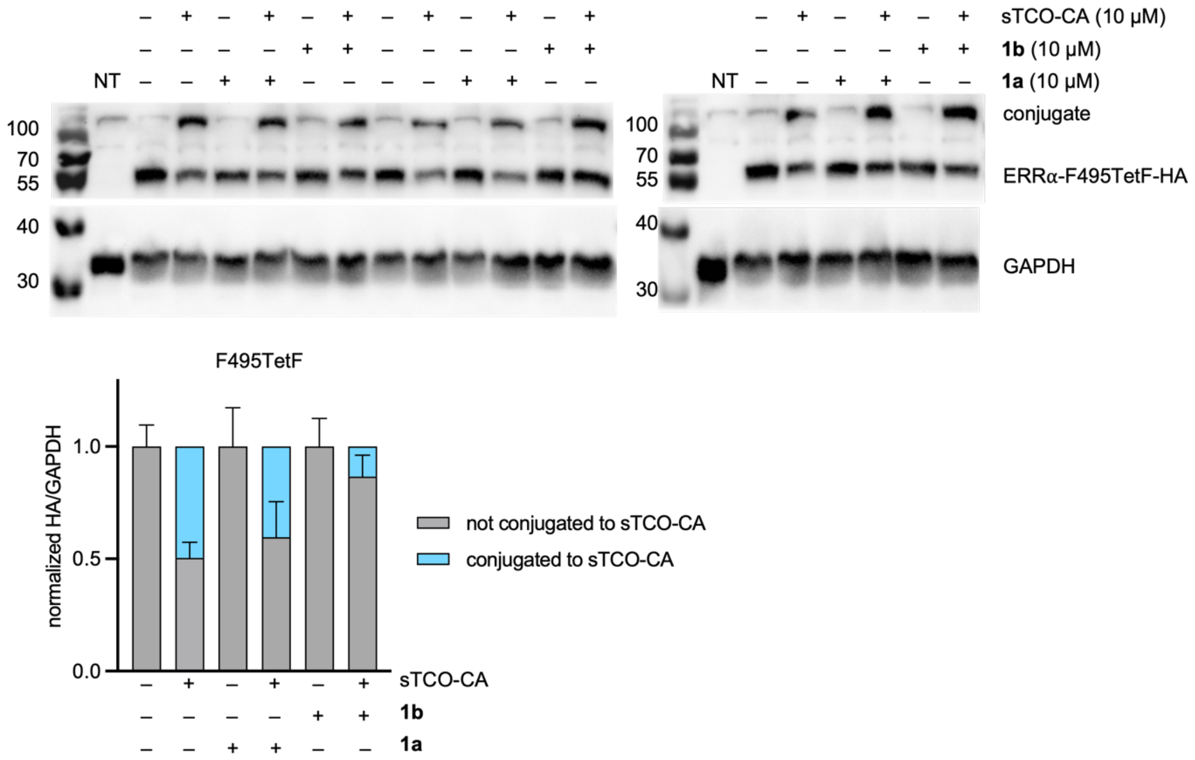

**Figure S19. Pulse-Chase Quantification of Bioconjugation for ERRα-F495TetF-HA.**

The bioconjugation efficiency of sTCO-CA, **1a**, and **1b** as determined via triplicate pulse-chase experiment in the presence of the proteasome inhibitor MG132 (10 µM).

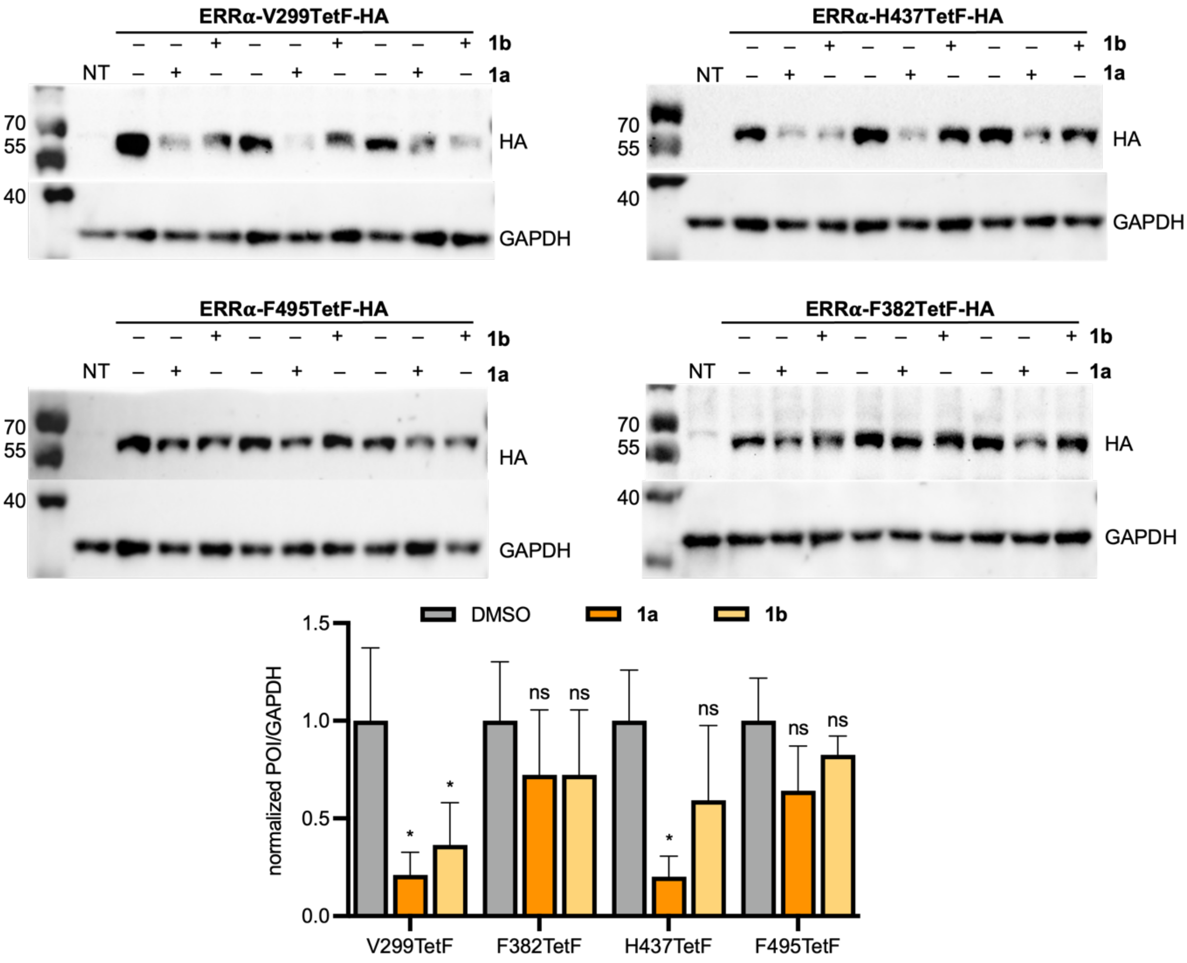

**Figure S20. Full HA and GAPDH blots for each quantification shown in Figure 5B.**

Error bars were calculated from three biological replicates, with statistical significance determined through one-way ANOVA or paired t-test, where * = $p < 0.1$, ** = $p < 0.01$, and *** = $p < 0.001$.

**Table S1. List of primers used for plasmid construction.**

Base mutations are indicated by capitalization.

| Primer | Sequence (5' to 3') | Annealing Temp (°C) | Elongation time (min) |
|---|---|---|---|
| P1 | cgcaaatgggcggtaggcgtg | - | - |
| P2 | gttcaggggggaggtgtg | - | - |
| P3 | ttcgaattcaccatggtgagcaagggcgaggagctgttcacc | 59.9 | 0.5 |
| P4 | ccttgctcaccttgtacagctcgtccatgccgccg | 59.9 | 0.5 |
| P5 | gctgtacaaggtgagcaagggcgaggagctgttcac | 60 | 3 |
| P6 | gcccttgctcaccatggtgaattcgaagcttgagctcgagatctgagtccggtag | 60 | 3 |
| P7 | caacgtcTAGatcatggccgacaagcagaagaacggcatcaag | 56.1 | 3.5 |
| P8 | gccatgatCTAgacgttgtggctgttgtagttgtactccagct | 56.1 | 3.5 |
| P9 | ctgaagggcTAGgacttcaaggaggacggcaacatcctgggg | 65.5 | 3.5 |
| P10 | cttgaagtcCTAgcccttcagctcgatgcggttcaccagggt | 65.5 | 3.5 |
| P11 | aaggaggacTAGaacatcctggggcacaagctggagtacaactacaac | 63.2 | 3.5 |
| P12 | caggatgttCTAgtcctccttgaagtcgatgcccttcagctcgat | 63.2 | 3.5 |
| P13 | ccagcagaacacccccatcggcgacTAGcccgt | 67 | 3.5 |
| P14 | ggttgtcgggcagcagcacgggCTAgtcgc | 67 | 3.5 |
| P15 | cgacaaccactacctgagcacccagtccTAGctgagcaa | 64 | 3.5 |
| P16 | gcgcttctcgttggggtctttgctcagCTAggactg | 64 | 3.5 |
| P17 | atccaccggtcgccacctacccatacgatgtt | 62.6 | 3 |
| P18 | tttaagcgtaatctggaacatcgtatgggtaggtggcgaccg | 62.6 | 3 |
| P19 | tacatcttgaacgtcacccccaatttgccgaatTAGtttgaga | 60.1 | 3 |
| P20 | atttgcttgtatttaaactctcctgcgttctcaaaCTAattcggca | 60.1 | 3 |
| P21 | gtttaaatacaagcaaatccccatctcggatcactggTAGcaaaacc | 61.1 | 3 |
| P22 | cctcagggaaaaactgggacaggttttgCTAccagtgat | 61.1 | 3 |
| P23 | aaaacctgtcccagtttttccctgaggccattTAGttcatagat | 61 | 3 |
| P24 | cttgccccgggcttcatctatgaaCTAaatggcctc | 61 | 3 |
| P25 | atgaaaaaatccaacatatcccctaacttcaacttcatgTAGcagctgc | 60.2 | 3 |
| P26 | gtcctctcgaagtccagcagctgCTAcatgaag | 60.2 | 3 |
| P27 | gtggcggccgccaccatggattacaaggatgacgacgataaggcccggg | 61.1 | 1.5 |
| P28 | atggtggcgtcccacccccttgcctgagtccatcatggc | 61.1 | 1.5 |
| P29 | ggtgggacgccaccatggtgagcaagggcgagga | 63.5 | 3 |
| P30 | atccttgtaatccatggtggcggccgccaccg | 63.5 | 3 |
| P31 | ggacgccacctacccatacgatgttccagattacgcttaaagcggccg | 61 | 3 |
| P32 | cgtatgggtaggtggcgtcccacccccttgc | 61 | 3 |
| P33 | tcccagccTAGgctaccctctgtgacctctttgaccgagagattgtggt | 66.2 | 3 |
| P34 | ggtagcCTAggctgggaggtgcccatcagggcctg | 66.2 | 3 |
| P35 | ctggccTAGgctgaggacttagtcctggatgaagaggggggca | 66 | 3 |
| P36 | tcctcagcCTAggccagctcatcctgcagtggcagtgagc | 66 | 3 |
| P37 | ctctgtgTAGatcgaagatgccgaggctgtg | 66 | 3 |
| P38 | cttcgatCTAcacagagtctgaattggcaagggc | 66 | 3 |
| P39 | gcccatTAGtatggggtgaagctggagggcaag | 62.2 | 3 |
| P40 | cccataCTAatgggccagcactttgcccgc | 62.2 | 3 |
| P41 | gcgcggcctcatcacaggttggtgctgat | 64 | 1 |
| P42 | aagctgtgaccggcgcctactatggcgtgtc | 64 | 1 |

### 4. $^1$H and $^{13}$C NMR Spectra

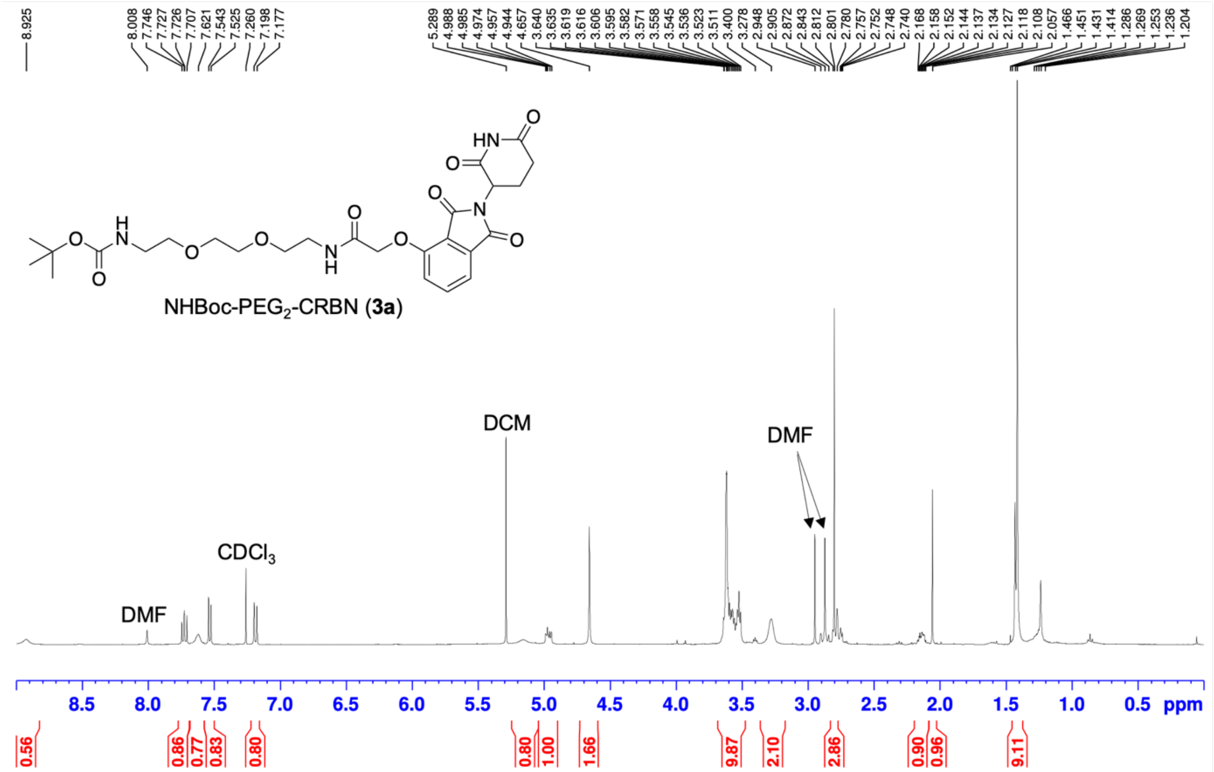

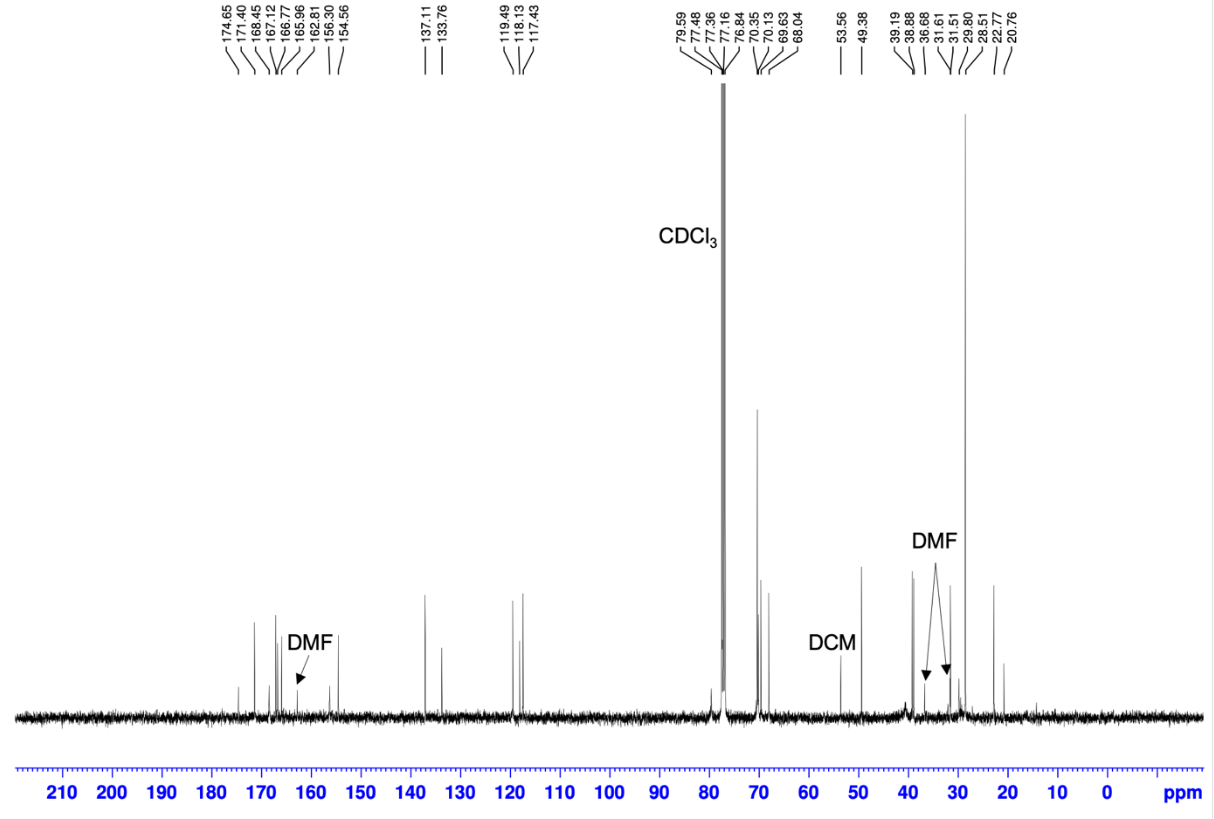

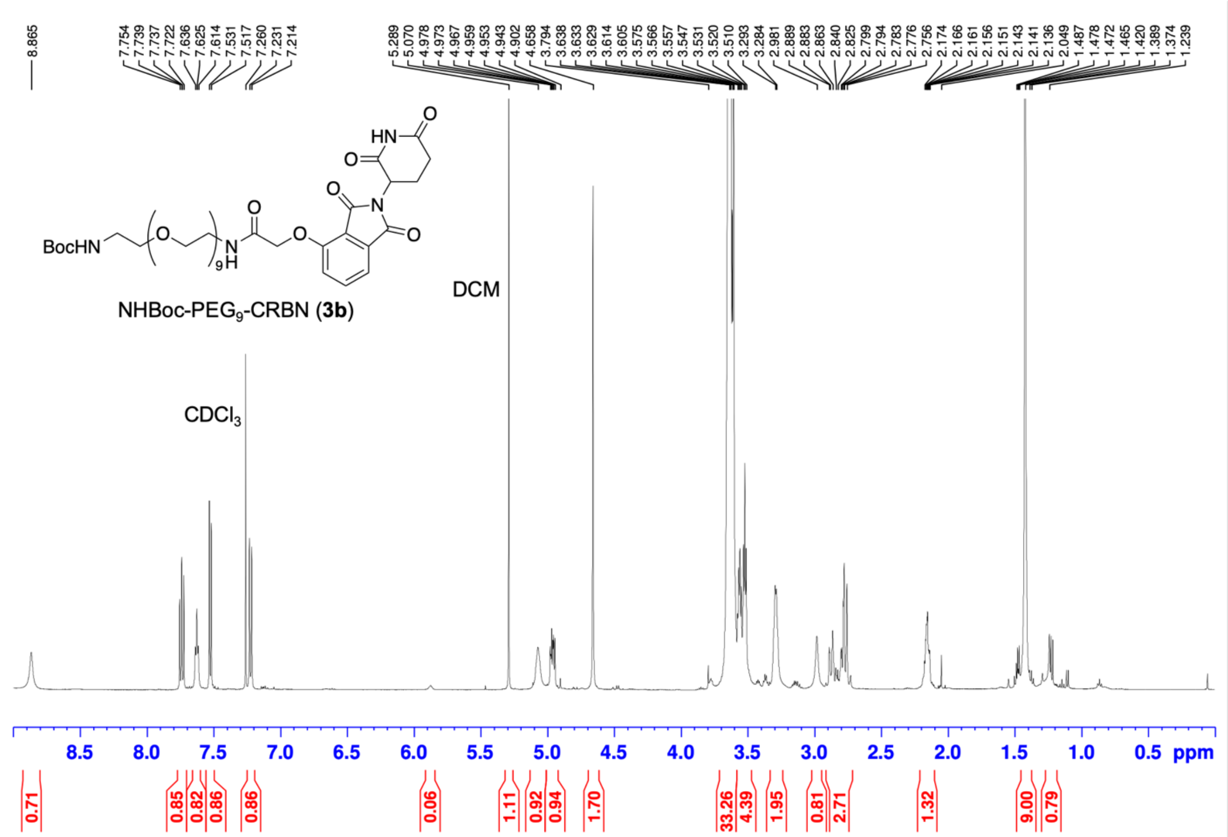
8.865
7.754 7.739 7.737 7.722 7.636 7.625 7.614 7.531 7.517 7.260 7.231 7.214
5.289 5.070 4.978 4.973 4.967 4.959 4.953 4.943 4.902 4.658 3.794 3.638 3.633 3.629 3.614 3.605 3.575 3.566 3.557 3.547 3.531 3.520 3.510 3.293 3.284 2.981 2.889 2.883 2.863 2.840 2.825 2.799 2.794 2.783 2.776 2.756 2.174 2.166 2.161 2.156 2.151 2.143 2.141 2.136 2.049 1.487 1.478 1.472 1.465 1.420 1.389 1.374 1.239
BocHN
O
N
H
9
HN
N
NHBoc-PEG9-CRBN (3b)
DCM
CDCl3
8.5 8.0 7.5 7.0 6.5 6.0 5.5 5.0 4.5 4.0 3.5 3.0 2.5 2.0 1.5 1.0 0.5 ppm
0.71 0.85 0.82 0.86 0.86 0.06 1.11 0.92 0.94 1.70 33.26 4.39 1.95 0.81 2.71 1.32 9.00 0.79

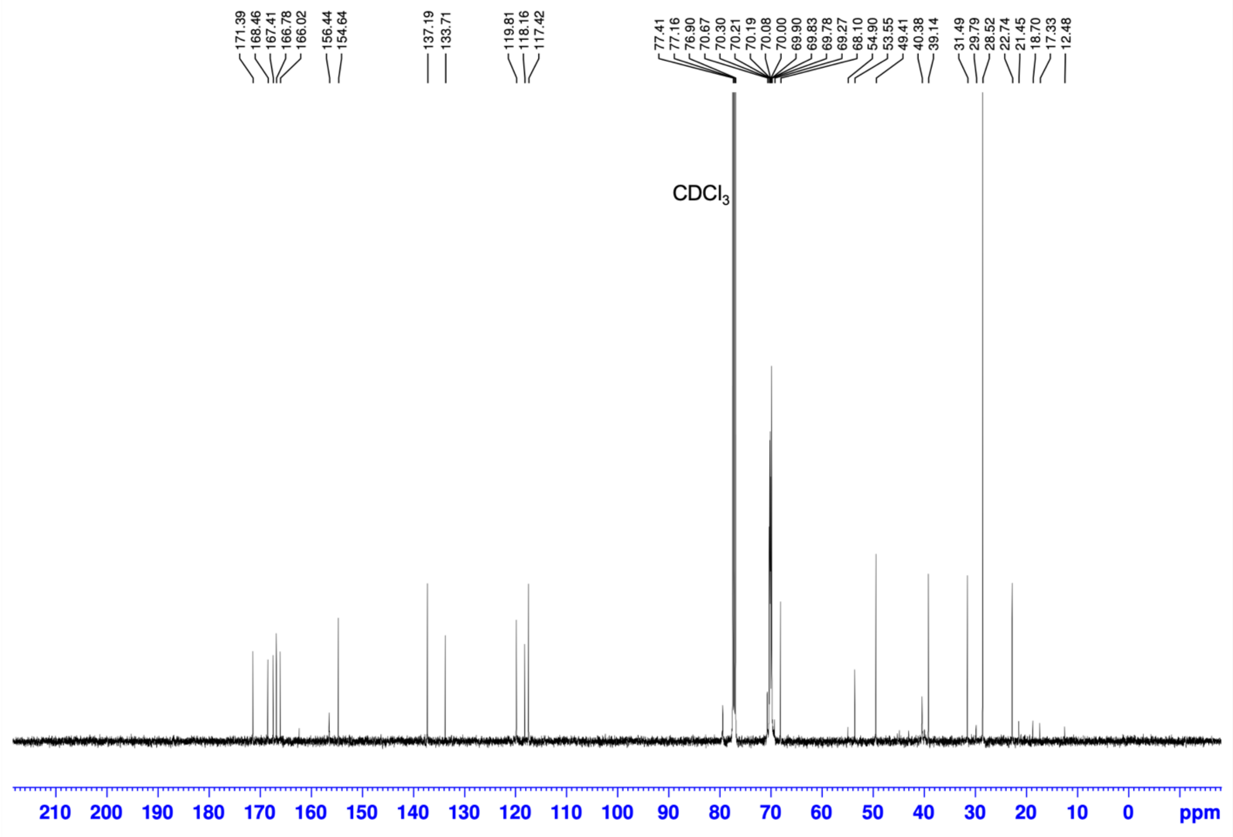
171.39 168.46 167.41 166.78 166.02 156.44 154.64 137.19 133.71 119.81 118.16 117.42
77.41 77.16 76.90 70.67 70.30 70.21 70.19 70.08 70.00 69.90 69.83 69.78 69.27 68.10 54.90 53.55 49.41 40.38 39.14 31.49 29.79 28.52 22.74 21.45 18.70 17.33 12.48
CDCl3
210 200 190 180 170 160 150 140 130 120 110 100 90 80 70 60 50 40 30 20 10 0 ppm

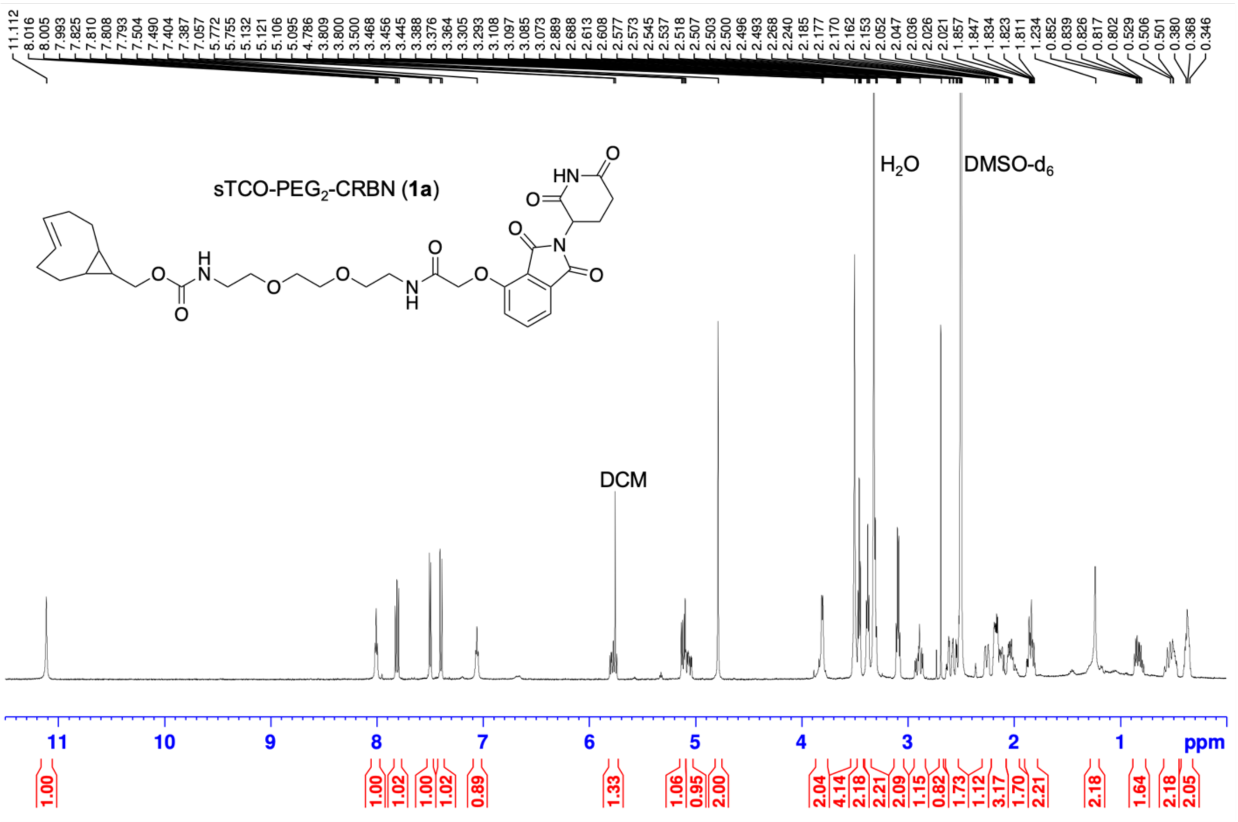
11.112
8.016
8.005
7.993
7.825
7.810
7.808
7.793
7.504
7.490
7.404
7.387
7.057
5.772
5.755
5.132
5.121
5.106
5.095
4.786
3.809
3.800
3.500
3.468
3.456
3.445
3.388
3.376
3.364
3.305
3.293
3.108
3.097
3.085
3.073
2.889
2.688
2.613
2.608
2.577
2.573
2.545
2.537
2.518
2.507
2.503
2.500
2.496
2.493
2.268
2.240
2.185
2.177
2.170
2.162
2.153
2.052
2.047
2.036
2.026
2.021
1.857
1.847
1.834
1.823
1.811
1.234
0.852
0.839
0.826
0.817
0.802
0.529
0.506
0.501
0.380
0.368
0.346
sTCO-PEG2-CRBN (1a)
HN
O
N
H
H2O
DMSO-d6
DCM
11
10
9
8
7
6
5
4
3
2
1
ppm
1.00
1.00
1.02
1.00
1.02
0.89
1.33
1.06
0.95
2.00
2.04
4.14
2.18
2.21
2.09
1.15
0.82
1.73
1.12
3.17
1.70
2.21
2.18
1.64
2.18
2.05

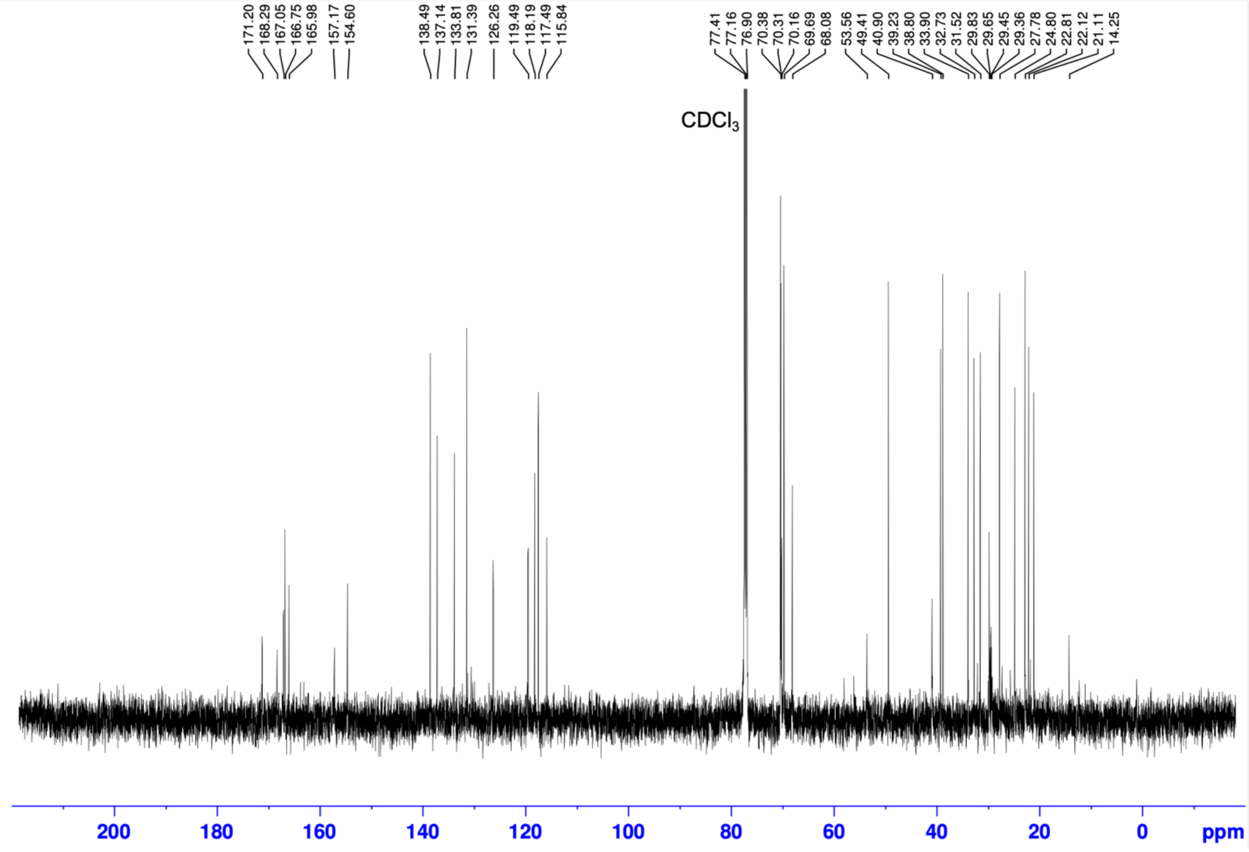
171.20
168.29
167.05
166.75
165.98
157.17
154.60
138.49
137.14
133.81
131.39
126.26
119.49
118.19
117.49
115.84
77.41
77.16
76.90
70.38
70.31
70.16
69.69
68.08
53.56
49.41
40.90
39.23
38.80
33.90
32.73
31.52
29.83
29.65
29.45
29.36
27.78
24.80
22.81
22.12
21.11
14.25
CDCl3
200
180
160
140
120
100
80
60
40
20
0
ppm

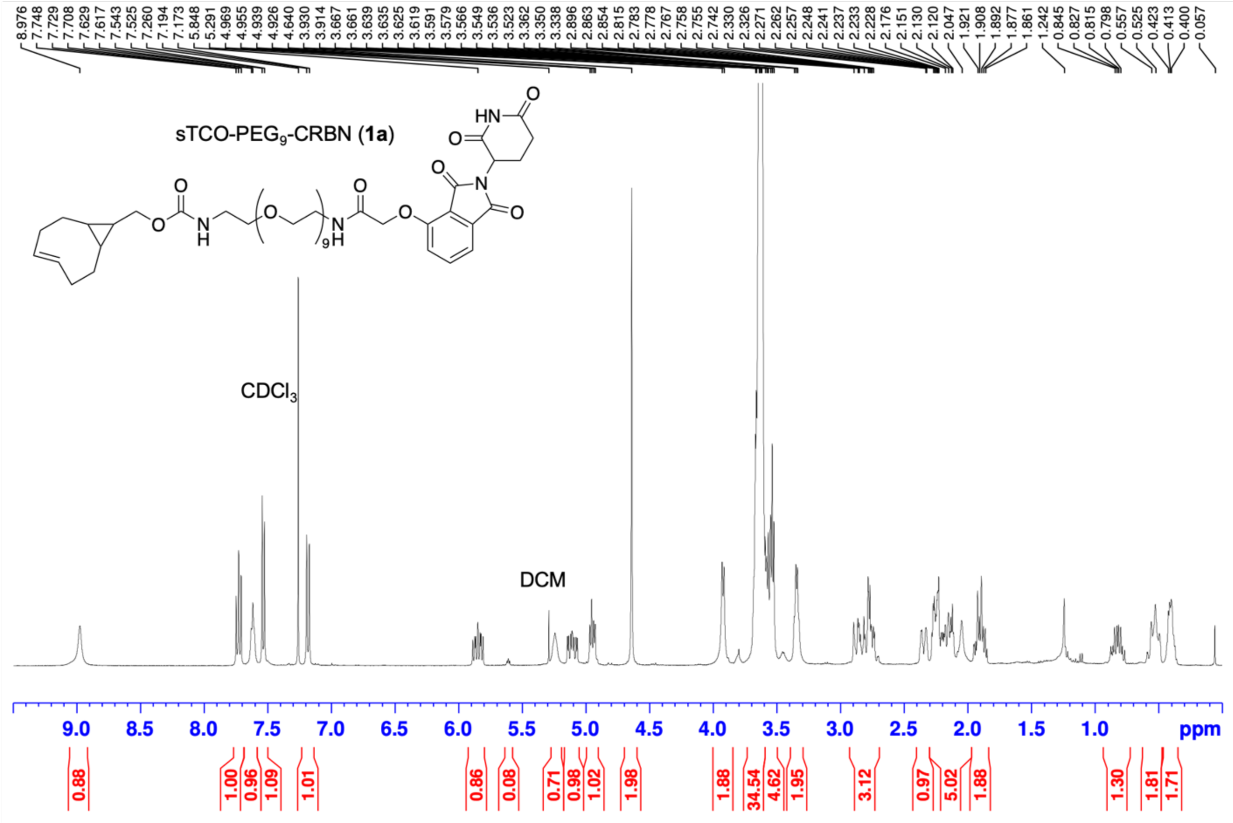
sTCO-PEG$_9$-CRBN (**1a**)
8.976 7.748 7.729 7.708 7.629 7.617 7.543 7.525 7.260 7.194 7.173 5.848 5.291 4.969 4.955 4.939 4.926 4.640 3.930 3.914 3.667 3.661 3.639 3.635 3.625 3.619 3.591 3.579 3.566 3.549 3.536 3.523 3.362 3.350 3.338 2.896 2.863 2.854 2.815 2.783 2.778 2.767 2.758 2.755 2.742 2.330 2.326 2.271 2.262 2.257 2.248 2.241 2.237 2.233 2.228 2.176 2.151 2.130 2.120 2.047 1.921 1.908 1.892 1.877 1.861 1.242 0.845 0.827 0.815 0.798 0.557 0.525 0.423 0.413 0.400 0.057
CDCl$_3$
DCM
9.0 8.5 8.0 7.5 7.0 6.5 6.0 5.5 5.0 4.5 4.0 3.5 3.0 2.5 2.0 1.5 1.0 ppm
0.88 1.00 0.96 1.09 1.01 0.86 0.08 0.71 0.98 1.02 1.98 1.88 34.54 4.62 1.95 3.12 0.97 5.02 1.88 1.30 1.81 1.71

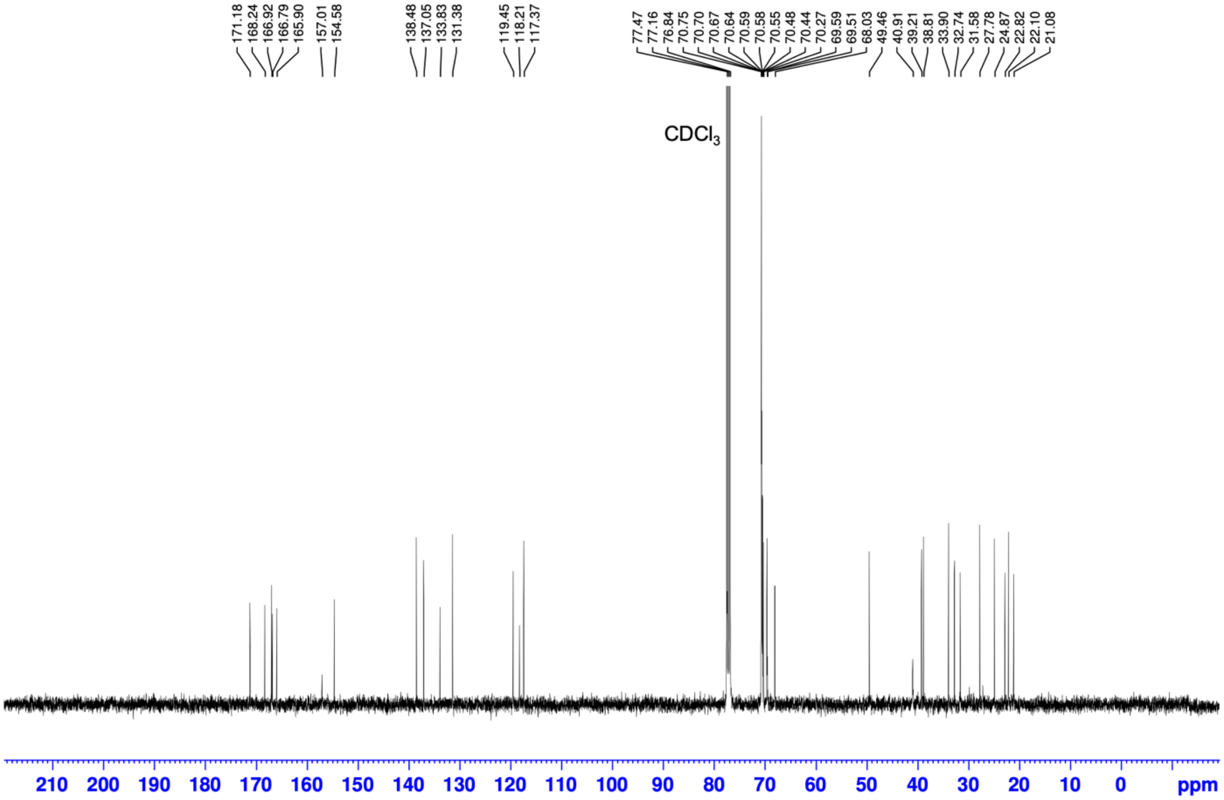
171.18 168.24 166.92 166.79 165.90 157.01 154.58 138.48 137.05 133.83 131.38 119.45 118.21 117.37 77.47 77.16 76.84 70.75 70.70 70.67 70.64 70.59 70.58 70.55 70.48 70.44 70.27 69.59 69.51 68.03 49.46 40.91 39.21 38.81 33.90 32.74 31.58 27.78 24.87 22.82 22.10 21.08
CDCl$_3$
210 200 190 180 170 160 150 140 130 120 110 100 90 80 70 60 50 40 30 20 10 0 ppm
